# Reversible Antagonism of Dopamine D1 Receptor using a Photoswitchable Remotely Tethered Ligand

**DOI:** 10.1101/2025.06.10.658926

**Authors:** Belinda E. Hetzler, Prashant Donthamsetti, Robert M. Wolesensky, Cherise Stanley, Ehud Y. Isacoff, Dirk Trauner

**Author notes:** Corresponding Author Dirk Trauner − Department of Chemistry, New York University, New York, New York 10003, United States; Department of Chemistry and Department of Systems Pharmacology and Translational Therapeutics, University of Pennsylvania, Philadelphia 19104, United States;, Ehud Y. Isacoff − Department of Molecular and Cell Biology and Helen Wills Neuroscience Institute, University of California, Berkeley, California 94720, United States; Molecular Biophysics & Integrated Bioimaging Division, Lawrence Berkeley National Laboratory, Berkeley, California 94720, United States;, Prashant Donthamsetti − Department of Pharmacology, Vanderbilt University, Nashville, Tennessee 37232, United States. These authors contributed equally (Belinda E. Hetzler, Prashant Donthamsetti).

## Abstract

Dopamine D1 receptor (D1R) plays key roles in health and disease. D1R is broadly expressed throughout the brain and body and is dynamically activated in response to endogenous dopamine, making it difficult to target this receptor with sufficient precision. We previously developed a robust light-activatable, tetherable agonist for D1R, wherein a temporally precise photo-switch (the P compound) binds to a genetically-encoded membrane anchoring protein (the M protein) in specific brain locations and cell types. Here we extended our approach by developing a complementary antagonist P compound that could be used to block specific populations of D1R in the brain with precise timing. Together, we have generated a robust toolkit for interrogating D1R function in the brain with unprecedented precision.

## Introduction

The neuromodulator dopamine play key roles in health (movement, reward, motivation, aversion, vasodilation) and disease (Parkinson’s disease, schizophrenia, addiction, hypertension) via its actions on five G protein-coupled dopamine receptors.^1–7^ Dopamine receptors are divided into two subfamilies, the G_s/olf_-coupled D1-like receptors (D1R, D5R) and the G_i/o/z_-coupled D2-like receptors (D2R, D3R, D4R). Considerable effort has gone towards developing small molecule dopamine receptor ligands for the treatment of disease. For example, D1-like receptors are putative targets for treatment of central nervous system (CNS) disorders such as Parkinson’s Disease, and they have been targeted peripherally for hypertension.^8–12^ However, D1-like receptors are broadly expressed throughout the body, including outside the nervous system, and are activated with complex temporal dynamics, making it difficult to identify precisely which receptor populations must be targeted to maximize therapeutic efficacy and avoided to minimize adverse side effects.

Existing methods for interrogating the function of D1-like receptors lack precision. Most conventional small molecule drugs cannot differentiate between D1R and D5R due to the high degree of homology in the orthosteric dopamine binding site.^13^ In addition, these agents are freely diffusible so cannot be used to target specific cell types and tissues. Genetic approaches (knockin, knockout, overexpression) are molecularly and spatially specific but are long lasting, which limits our understanding of the temporal dynamics of receptor activation and are prone to compensatory changes in gene expression. Chemogenetic (DREADDs) and optogenetic (opto-XRs) techniques can be used to engage signaling with cellular and spatiotemporal specificity, but these tools are ectopically overexpressed unnatural proteins that lack biochemical selectivity for dopamine and potentially overdrive the system.^14,15^

We have had a long-standing interest in developing chemical tools to interrogate complex receptor systems and deconvolve the contribution of individual receptor subtypes in specific cells to neural circuit function.^16,17^ In an approach called photo-pharmacology, a photo-chromic ligand can be rendered biologically active or inactive under the action of light.^18^ In its simplest conception, a photo-reactive protecting group can be removed under irradiation with light to reveal the biologically active ligand that remains active until it is removed by diffusion. For example, photo-caged dopamine can enable timed dopamine release by light.^19–21^

Alternatively, a ligand can also be equipped with a permanent photo-chromic unit that is incorporated into its scaffold. For example, azobenzene-based photo-switches can be reversibly toggled between a *cis*- and *trans*-configuration, with one form ideally being biologically inactive and the other being active.^18^ However, these compounds diffuse out of the illuminated target area, undermining spatial precision. Moreover, while light can provide spatial focus to tissues and tissue areas of interest for switching the ligand to an active or available state, dopamine receptors are expressed in different cell types within the same brain area.

We recently solved the problems of diffusion and cell-specificity using an approach called membrane-anchored photo-switchable orthogonal remotely tethered ligand (MP), which has two components (**Fig. 1**).^22–24^ The first component is a photo-switchable ligand (the P compound) that is linked to a reactive substrate (e.g., benzylguanine) via a flexible linker (**Fig. 1A**). The second component is a genetically-encoded membrane-anchoring protein (the M protein) that contains a self-labeling protein-tag (e.g., SNAP-tag) that is compatible with the reactive substrate, which is selectively expressed in the cell type of interest (**Fig. 1B**). After the P compound attaches to the M protein in a chemically specific covalent labeling event, the MP complex redistributes in the cell membrane, where it encounters the target receptor, thereby placing the P within reach of the receptor ligand binding site so that the ligand is able to bind in a light-gated manner (**Fig. 1B**).

**Figure 1.**
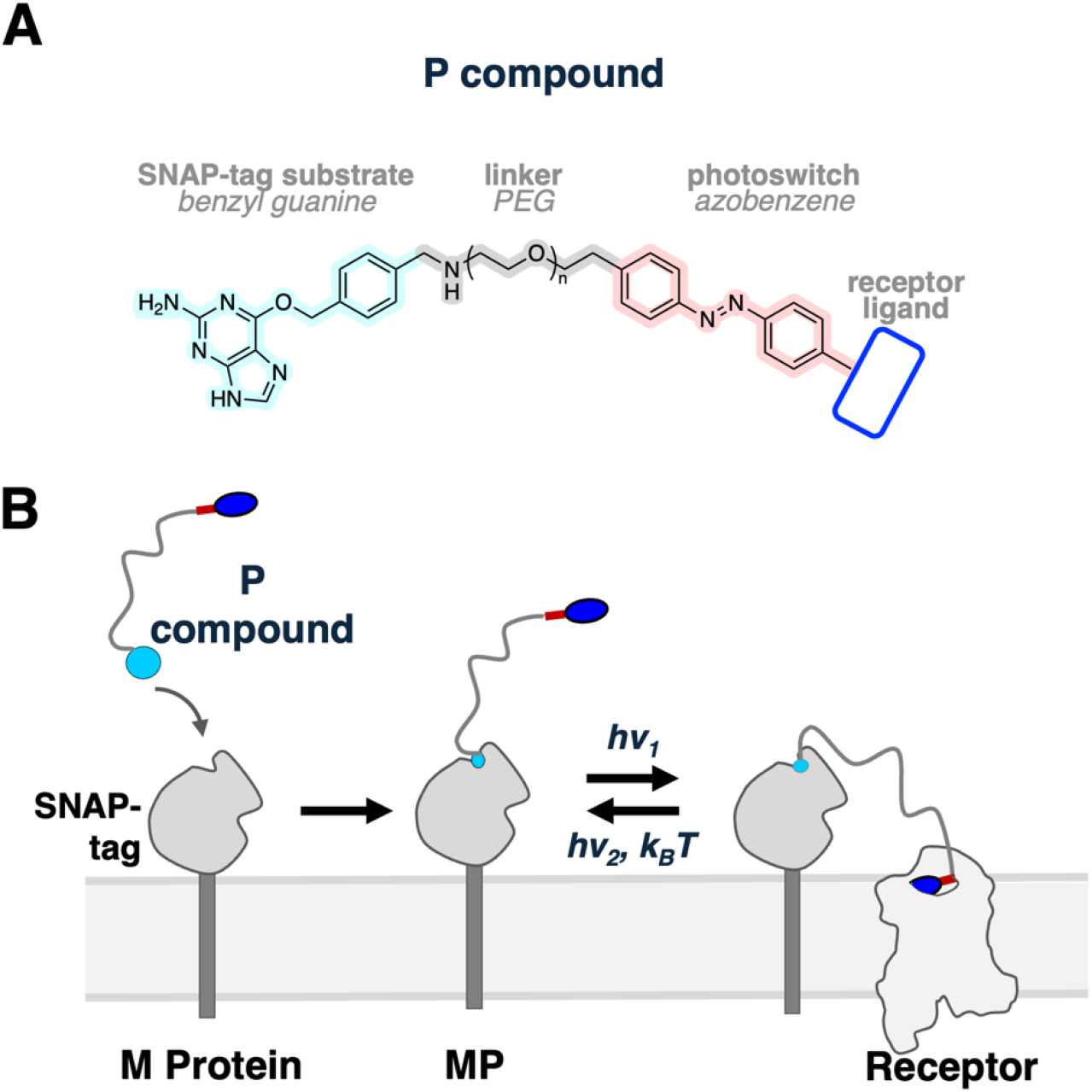
Design of a photo-pharmacological system to control a GPCR such as D1R in a cell type-specific and spatiotemporal manner. (**A**) The chemical component (the P compound) is comprised of a SNAP-tag reactive benzylguanine linked via a flexible polyethylene (PEG)-linker to a photo-switchable ligand. (**B**) Specific cells are genetically modified to express the membrane anchored protein (M), a SNAP-tag anchored to the cell membrane via a single-pass transmembrane segment anchor. After application of the P compound to M protein-expressing cells, the covalent MP complex is formed. The P reaches from the M to the receptor and reversibly binds in response to a specific wavelength of light.

We recently developed D2-like receptor selective agonist and antagonist MPs that are tethered via chloroalkane to a HaloTag M protein (**MP-D2**_**ago**_ and **MP-D2**_**block**_, respectively).^24^ We also developed a D1-like receptor agonist MP that can be tethered via benzylguanine to a SNAP-tagged M protein.^16,23^ This compound, **MP-D1**_**ago**_, is based on the ligand PPHT and operates as a bistable photo-switch that is active in the *trans-*configuration under blue light and inactive in the *cis-*configuration under near-UV light. Using this *trans*-active **MP-D1**_**ago**_ system, we found that the acute activation of D1R in the dorsal striatum of mice promotes movement initiation and reinforces antecedent patterns of cortical brain activity.^16,25^

In this study, we set out to develop a light-mediated D1-like receptor antagonist MP to complement our existing dopamine receptor MP toolkit. We designed compounds that conjugate to SNAP-tag, enabling future multiplexed experiments wherein SNAP-M for D1-like receptors and Halo-M for D2-like receptors can be controlled independently. Here, we report the development of a thermally bistable D1R receptor-specific tethered antagonist (**MP-D1**_**block**_) that completes a set of tethered ligands to activate or inactivate dopamine receptors.^16,23,24^ This new MP will provide the chemical basis to study cell-specific D1R action *in vivo*.

## Results and Discussion

D1R can been blocked by synthetic small molecule inhibitors based on a tetrahydro-1H-3-benzazepine scaffold (**Fig. 2**). One of the first reported D1-like selective partial agonists was the catechol benzazepine **SKF38393** (**Fig. 2A**).^26–28^ With this scaffold, the biologically active conformation of dopamine is mimicked by constraining the amine into a seven-membered ring.^29^ The substitution pattern on the aryl moiety was subsequently optimized, leading to the discovery of the D1 partial agonist drug **fenoldopam** (**Fig. 2A**), which has an additional chlorine substituent and a phenol instead of a phenyl group.^30^ Within these benzazepine derivatives, the benzylic stereocenter is important, with the *S*-enantiomer being virtually inactive.^31–33^ The exchange of one catechol hydroxy group to a halogen and *N*-Methylation led to the development of D1-like receptor selective antagonists such as aryl-bromide variant **SKF83566**.^34^ Further modifications on the benzazepine scaffold include the rigidification of the pendant phenyl with a ethane bridge onto the azepine moiety, yielding D1R antagonist **ecopipam** (**Fig. 2A**), which is currently under clinical investigations for several CNS disorders.^12^

**Figure 2.**
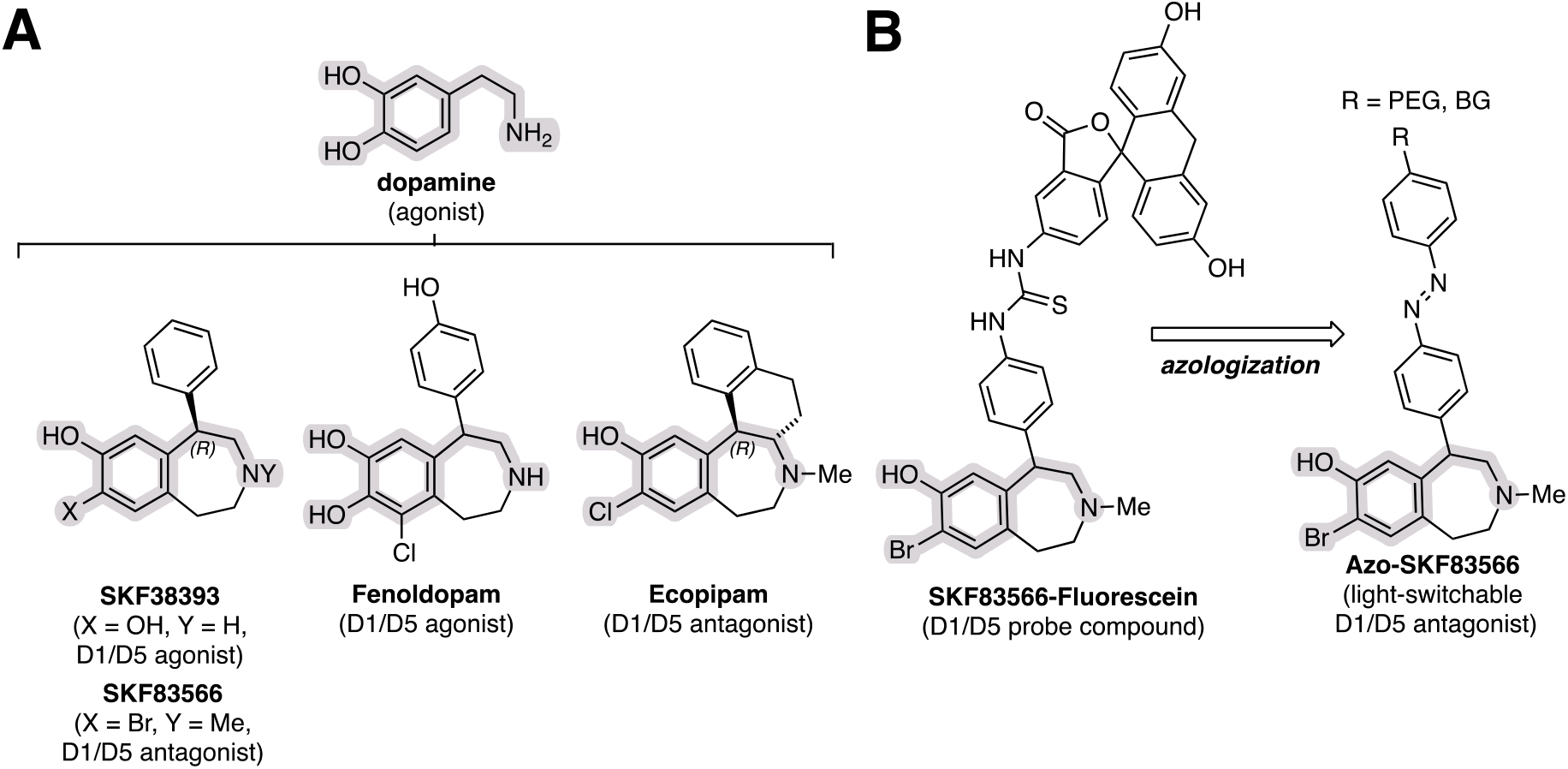
Design of a D1R MP blocker. (**A**) Dopamine and conformationally restricted D1R partial agonists and antagonists sharing a common tetrahydro-1H-3-benzazepine scaffold. (**B**) Tool compounds based on D1R antagonist **SKF83566** (such as **SKF83566-Fluorescein**) indicate that the elongation of the phenyl ring in the *para*-position is tolerated without compromising potency. This site is suitable for the incorporation of azobenzene connected to a PEG linker and benzylguanine (BG) at that position for the design of a D1R tool for proximity photo-pharmacology.

A series of tetrahydro-1H-3-benzazepine based-D1R tool compounds were previously developed to visualize and study dopamine receptors in cells. Here, the appendant phenyl group on the tetrahydro-benzazepine was elongated with a short linker to connect to large residues, such as fluorescein and BODIPY (**Fig. 2B**) without compromising their ability to bind D1R.^35–37^ We identified the phenyl group as ideal azologization motif for the development of D1R antagonist P compound, as the phenyl can be extended with an azobenzene and linked to the PEG linker to the benzylguanine, the substrate for the SNAP-tag in the M protein.^38^ Recent cryoEM structures of a D1R-G_s_ complex bound to benzazepine agonists confirm that the phenyl lies at the solvent accessible top region of the orthosteric site in the seven transmembrane domain of the D1R, further supporting this proposed design of a D1R antagonist P compound.^39^

### Synthesis of P-D1_block12_ and P-D1_block24_

To set the stage for the installment of the azobenzene unit, we first prepared the known aniline **8** following literature reported methods(**Fig. 3A**):^35^ To this end, commercial bromoketone **1** was converted to epoxide **2** in excellent yield by reduction with NaBH_4_, followed by treatment with base. Epoxide opening of **2** with 4-methoxyphenylethylamine installed racemic phenylethanolamine derivative **4** that was cyclized to tetrahydro-1H-3-benzazepine **5** using polyphosphoric acid at 100 °C. In an Eschweiler–Clarke reaction, methylation of the secondary amine **5** was achieved in good yield. Resolution of **6*-rac*** using a chiral tartaric acid derivative yielded the biologically active *R*-enantiomer **6** in >99% *ee*, as determined by chiral HPLC. The sign of the optical rotation matched literature reported values, and absolute configuration was assumed based on literature precedence.^40^

**Figure 3.**
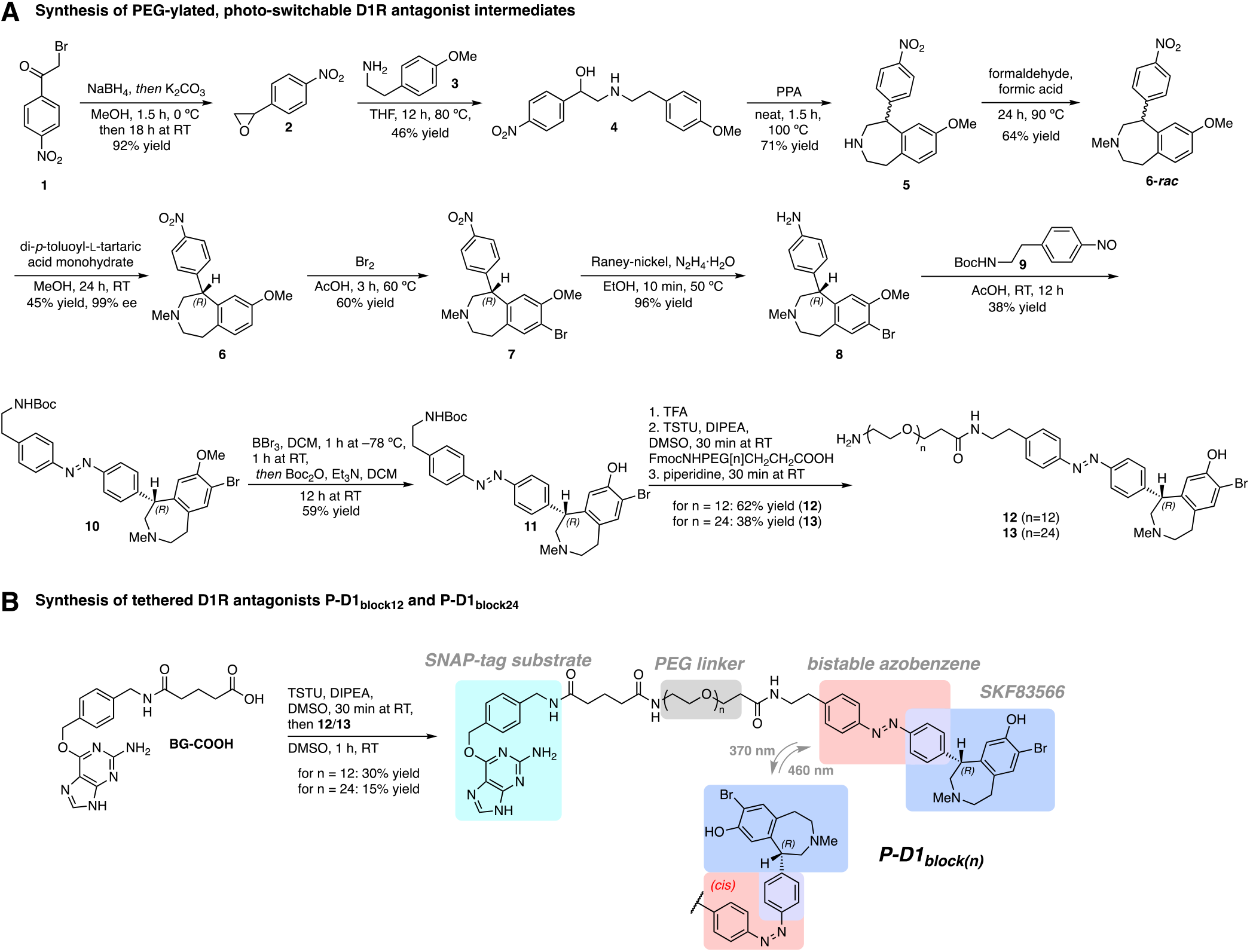
Multistep chemical synthesis of P-D1_block12_ and P-D1_block24_. (**A)** Synthesis of PEGylated D1R antagonist intermediates 12 and **13** harboring a photoswitchable azobenzene moiety. (**B)** Assembly of the tool compounds **P-D1**_**block12**_ and **P-D1**_**block24**_ by connecting intermediates **12** and **13** and to benzyl guanine carboxylic acid (**BG-COOH**).

Bromination followed by Raney-Nickel reduction furnished the desired aniline **8** in overall good yields that underwent, Baeyer– Mills reaction with freshly prepared nitroso-phenylethyl Boc-amine **9** in acceptable yields. Boron tribromide effected methoxy deprotection of **10**, but also resulted in Boc-deprotection. For purification purposes, we -protected the primary amine with Boc_2_O to yield **11** that was stable and could be purified on SiO_2_. Boc-deprotection of **11** with TFA provided a primary amine as the connecting point for introduction of the long flexible linker.

For our tethered photo-pharmacological approach, PEG-linkers have proven to be suitable as they are soluble, monodisperse, commercially available in different lengths, and biocompatible. Reaction of the crude primary amine in a TSTU-mediated peptide coupling with FmocNHPEG[n]CH_2_CH_2_COOH yielded photo-switchable D1R antagonists with the desired flexible linker. We chose to use PEG[12] and PEG[24], lengths that have previously successfully spanned the distance between orthosteric GPCR binding site and proximal M protein-bearing reactive tag.^17,24^ Piperidine effected Fmoc-deprotection to unveil primary amines **12** (PEG[12]) and **13** (PEG[24]) that were then coupled to the SNAP-tag substrate benzyl guanine (**BG-COOH**) by TSTU-mediated peptide coupling (**Fig. 3B**).^41^ Finally, the desired remotely tethered D1R antagonists **(*R*)*-*P-D1**_**block12**_ and **(*R*)*-*P-D1**_**block24**_ were obtained. The procedure starting from **6** was performed using pure (*R*)-intermediate. Additionally, an (*S*)-configured sample of **6** with 56% *ee*

### Photo-physical Characterization of P-D1_block(n)_

We then characterized the photo-physical properties of **P-D1**_**block24**_ by UV-Vis and NMR spectroscopy to confirm classic azobenzene behavior in response to light (**Fig. 4**). **P-D1**_**block12**_ was assumed to perform on par to **P-D1**_**block24**,_ as no change in the chromophore unit has occurred. Full thermally relaxed *trans*-**P-D1**_**block24**_ has an absorption maximum of 338 nm in DMSO (**Fig. 4A**). After irradiation with 370 nm light for 10 min, *cis*-enriched **P-D1**_**block24**_ absorbs visible light maximally at 436 nm and can be switched back into its *trans*-enriched form under irradiation with 460 nm light (10 min). **P-D1**_**block24**_ can be switched reversibly without fatigue using alternating 370 nm/460 nm light irradiation (**Fig. 4B**). We determined the photo-stationary state by pre-irradiating a 20 µM solution of **P-D1**_**block12**_ in DMSO at ambient temperature for 10 min and physically separating the sample into its *cis*- and *trans*-**P-D1**_**block24**_ isomers by LCMS to measure their ratio by peak integration. Under 360 nm, 25% *trans*-isomer remains, at 420 nm, 9% *cis*-isomer remains (**Fig. 4C**). The thermal relaxation half-life of the *cis*-isomer was determined in DMSO and 10% DMSO in PBS to confirm the bistable nature of the tool compound with a τ of 100.2 h and 11.8 h, respectively (**Fig. 4D**).

**Figure 4.**
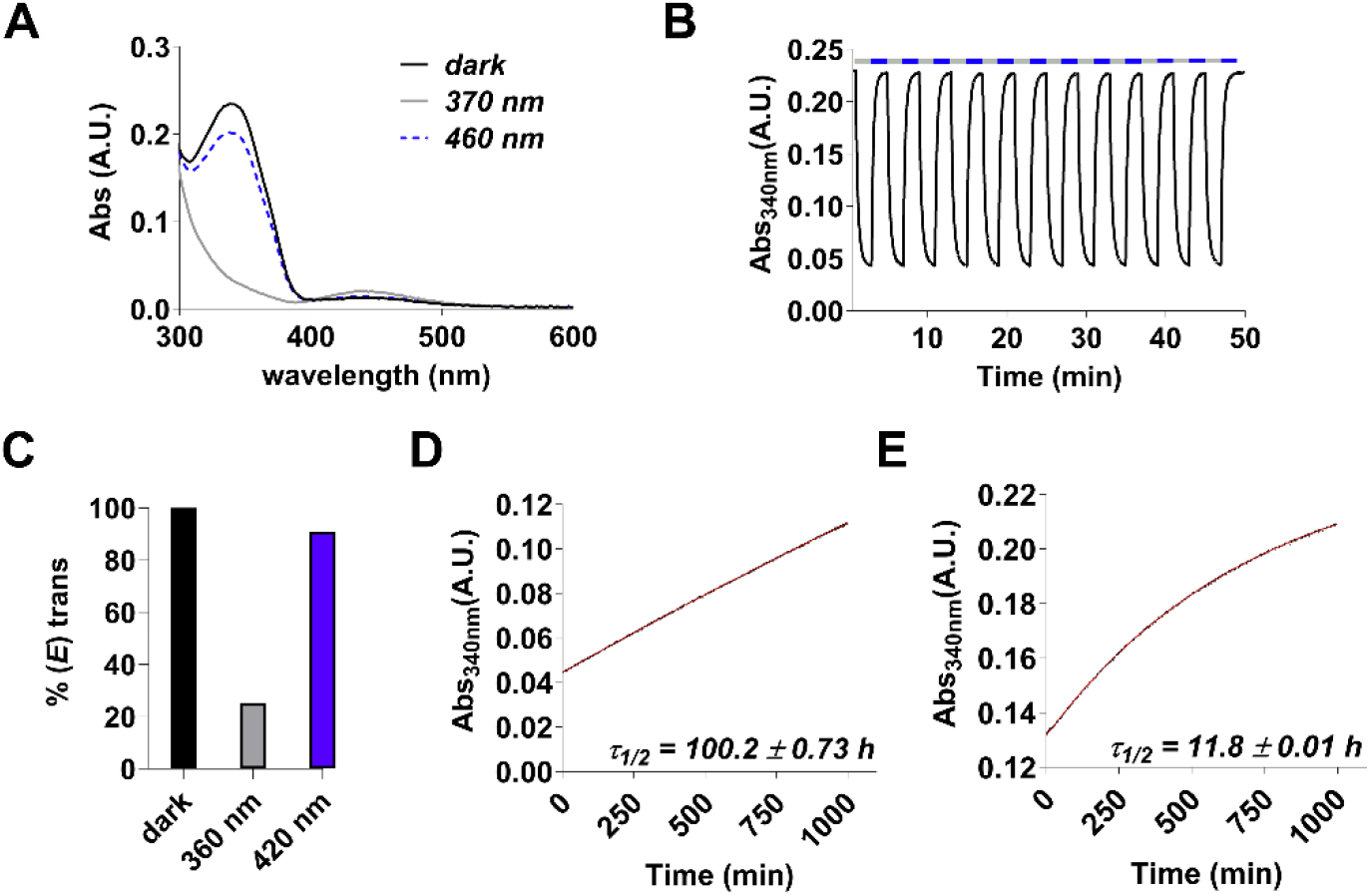
Photo-physical characterization of P-D1_block24_. **(A)** UV-Vis absorption spectra of dark-adapted (100% *trans*) **P-D1**_**block24**_ in DMSO (20 µM), and after pre-irradiation with 370/460 nm for 10 min. (**B)** Reversible switching of **P-D1**_**block24**_ without fatigue in DMSO (20 µM) under 370/460 nm irradiation. (**C)** Photo-stationary state (PSS) of **P-D1**_**block24**_ in DMSO (20 µM, 24 °C), after pre-irradiation with 360/420 nm for 10 min. Determination of the photo-stationary state was achieved by LCMS by separation of isomers and integration of the Abs at the isosbestic point of *cis*-and *trans*-isomers at the respective elution solvent mixtures. (**D, E) P-D1**_**block24**_ was determined to be thermally stable in 10% DMSO (*t*_1/2_ = 100 h) and 100% DMSO in PBS (*t*_1/2_ = 11.8 h) at 37 °C, detection of the Abs was performed at 340 nm after pre-irradiation of the samples for 10 min at 360 nm.

### Functional Analysis of MP-D1_block_

Next, we measured the effect on D1R of the **P-D1**_**block(n)**_ compounds when tethered to the M protein (MP-D1_block_). To determine D1R function, we used a whole-cell patch clamp electrophysiology assay in HEK293T cells that employs the G protein-activated inwardly rectifying K^+^ (GIRK) channel as an effector (**Supplementary Fig. 1**). Cells were co-expressed with D1R, GIRK, a modified G protein Gαi_s13_ that allows D1R to couple to GIRK, and the M protein (**Supplementary Fig. 1**). Following labeling with a **P-D1**_**block(n)**_ compound, cells were exposed to alternating 460 nm light and 370 nm light in the absence and presence of dopamine, which activates D1R and promotes an inward current via GIRK.

We first tested the effect of **(*S*)-P-D1**_**block12**_ tethered to the M protein on D1R. Dopamine-induced current decreased slightly in response to 370 nm light, an effect that was reversed in response to 460 nm light (**Fig. 5A,D**). In contrast, when tethered to the M protein, its enantiomer **(*R*)-P-D1**_**block12**_ robustly inhibited dopamine-induced current (**Fig. 5B,D**). Together, these results indicate that that **(*R*)-P-D1**_**block12**_ tethered to the M protein is a robust photo-antagonist of D1R in its *cis-*configuration and is inactive in its *trans*-configuration. The ability of **(*S*)-P-D1**_**block12**_ to weakly inhibit D1R is likely due to the presence of **(*R*)-P-D1**_**block12**_ in the sample (56% *ee*). Overall, this is consistent with earlier evidence that (*R*)-tetrahydro-1H-3-benzazepines being better binders of D1R than their (*S*)-enantiomers.^31,32^ Next, we tested the effect of extending the PEG linker from 12 to 24 PEGs. We observed **(*R*)-P-D1**_**block24**_ to have enhanced photo-block (**Fig. 5C,D**), with the potency of dopamine reduced by ∼216-fold in the *cis*-configuration (**Fig. 5E**).

**Figure 5.**
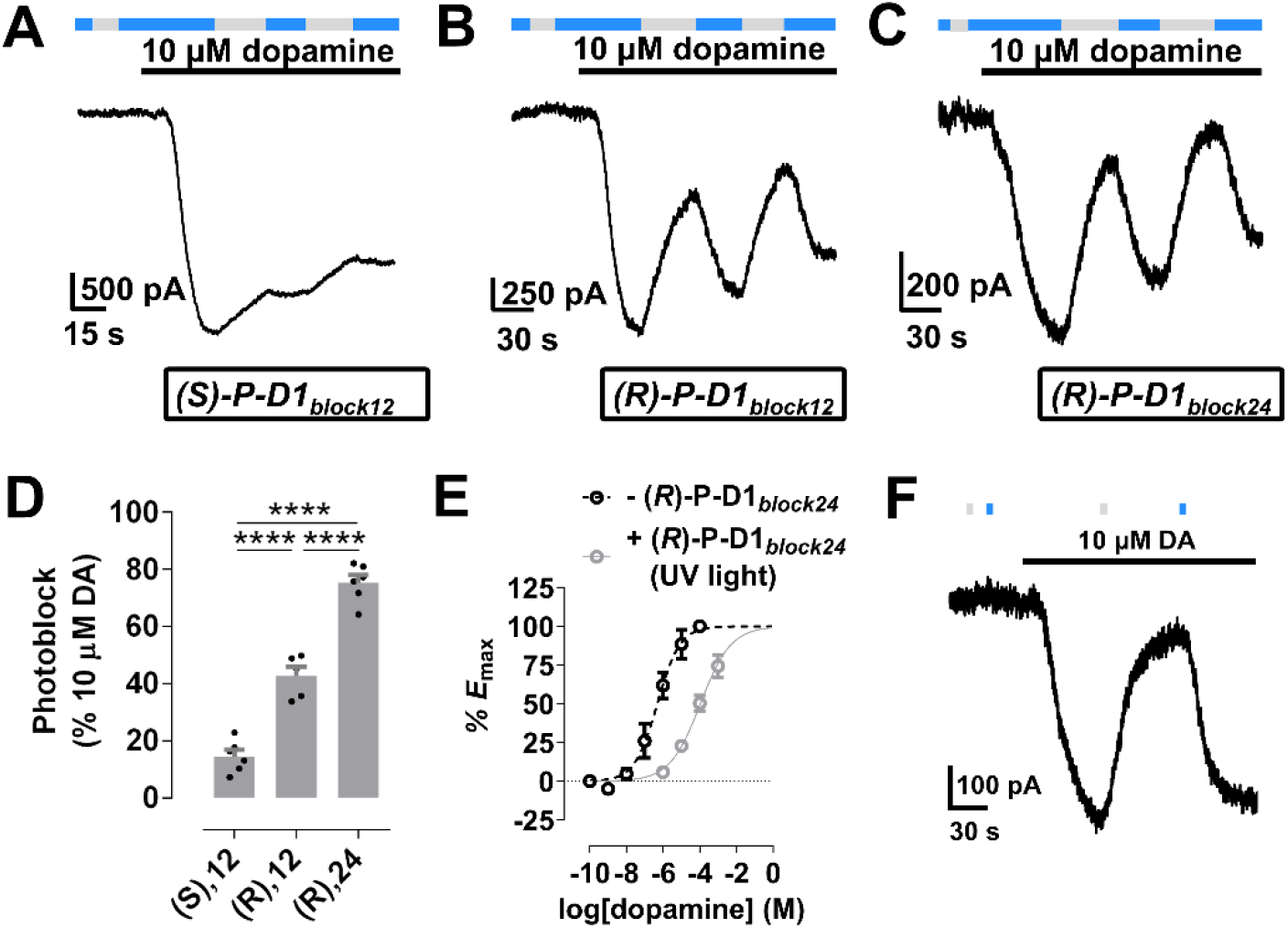
MP-D1_block_ photo-blocks D1R. (**A**) When tethered to the M protein, **(S)-P-D1**_**block12**_ weakly blocks D1R in its *cis*-configuration, which is likely due to the presence of the (*R*)-configured enantiomer in the sample (56% ee; 22% (*R*)-content). (**B-D) (*R*)-P-D1**_**block24**_ tethered to the M protein more effectively inhibits D1R than **(*S*)-P-D1**_**block12**_ or **(*R*)-P-D1**_**block12**_ tethered to the M protein in response to 370 nm light. (**E**) **(*R*)-P-D1**_**block24**_ tethered to the M protein decreases the potency of dopamine at D1R, consistent with competitive antagonism. (**F**) Consistent with its bistability, **(*R*)-P-D1**_**block24**_ tethered to the M protein elicits sustained inhibition of D1R in the dark following a short pulse of 370 nm light (grey bar), a process that is reversed with a short pulse of 460 nm light (blue bar).

Consistent with the thermal stability of **(*R*)-P-D1**_**block24**_, only a short pulse of light was required to inhibit (370 nm) or disinhibit (460 nm) dopamine-induced receptor activation (**Fig. 5F**).

## Conclusion

Together with our previous work, we have successfully completed a robust chemical toolbox of photo-switchable agonist and antagonist P compounds that can rapidly activate or prevent the activation of endogenous D1R and D2R.^16,24^ By engineering a photo-chromic azobenzene unit into a dopamine receptor ligand, a specific wavelength of light toggles the P compound into a bioactive configuration that binds to the receptor orthosteric site and a different wavelength of light that reverses the compound back to its binding-incompatible inactive configuration. The P compound is restricted to a desired cell type and location via covalent attachment to a genetically-encoded membrane-anchored self-labeling protein tag (the M protein). On the membrane of that cell, the MP complex interacts with the target receptor via random collision on the cell surface. Here, we report the development of a novel light-controllable, reversible, and cell type-specific D1R antagonist (**MP-D1**_**block**_) based on a tetrahydro-1H-3-benzazepine scaffold.

We have previously developed a robust D1R MP agonist (**MP-D1**_**ago**_) and demonstrated that its photo-activation in striatal direct-pathway medium spiny neurons (dMSNs) increases movement initiation and reinforces anteceding cortical patterns. ^16,25^ This agonist tool enables selective and precisely timed activation of D1R in living animals and is therefore able to reveal whether a receptor in a particular cell is *sufficient* for a specific behavior. In parallel, **MP-D1**_**block**_ could be used to block endogenous dopamine from activating D1R and to, thereby, reveal whether the receptor is *necessary* for the behavior. Thus, the combination of **MP-D1**_**ago**_ and **MP-D1**_**block**_ could provide a comprehensive view of the functional role of D1R in the brain.

When released, dopamine co-activates multiple dopamine receptors at once. Based on conventional behavioral pharmacology studies in rodents, dopamine receptor co-activation has been suggested to synergistically control downstream behaviors such as movement.^42,43^ However, because conventional D1R agonists and antagonists bind this receptor throughout the body, it is unclear which dopamine receptor populations in which cell types drive these effects. By employing orthogonal SNAP-tag labeling for our D1R MPs and HaloTag labeling for our D2R MPs, we can now selectively and simultaneously target multiple dopamine receptors, offering unique insight into dopamine action in the brain.

D1R is an attractive but historically intractable target for treating Parkinson’s Disease and other CNS disorders. This is in part because existing drug scaffolds have poor physiochemical properties, but also because these drugs broadly affect D1Rs across the brain and body. In addition, existing methods lack cellular and spatiotemporal precision, hindering identification the D1R populations that need to be targeted to treat disease. Our D1R MPs offer a unique opportunity to systematically target specific D1Rs in the brain and periphery and uncover their clinical potential. Furthermore, MP and related technologies could potentially be used in the clinic to treat disease with unprecedented precision, maximizing therapeutic efficacy and minimizing off-target effects.

## ASSOCIATED CONTENT

Supporting Information

The Supporting Information can be found on the bioRxiv website.

### Notes

The authors declare no competing financial interests.

## ACKNOWLEDGMENT

This work was supported by the National Institutes of Health (R01NS108151, D.T.; RF1MH123246, E.Y.I. and D.T.), the McKnight Endowment for Neuroscience (048240; E.Y.I. and D.T.), the European Research Council (268795; D.T.), Weill Neurohub (E.Y.I.), and the Brain and Behavior Research Foundation (P.D.). B.E.H. was supported by a MacCracken Fellowship. Robert Wolesensky thanks the Dean’s Undergraduate Research Fund at New York University for support. We thank New York University for financial support. NMR spectra were acquired on a Bruker Avance III 600 MHz using a TCI cryoprobe supported by the National Institutes of Health (S10 Award OD016343).

## Notes

### Competing Interest Statement

The authors have declared no competing interest.

## REFERENCES

(1) Beaulieu, J.-M.; Gainetdinov, R. R. The Physiology, Signaling, and Pharmacology of Dopamine Receptors. Pharmacol Rev 2011, 63 (1), 182–217. 10.1124/pr.110.002642.

(2) Beaulieu, J.-M.; Espinoza, S.; Gainetdinov, R. R. Dopamine Receptors – IUPHAR Review 13. Br. J. Pharmacol. 2015, 172 (1), 1–23. 10.1111/bph.12906.

(3) Klein, M. O.; Battagello, D. S.; Cardoso, A. R.; Hauser, D. N.; Bittencourt, J. C.; Correa, R. G. Dopamine: Functions, Signaling, and Association with Neurological Diseases. Cell Mol Neurobiol 2019, 39 (1), 31–59. 10.1007/s10571-018-0632-3.

(4) Martel, J. C.; Gatti McArthur, S. Dopamine Receptor Subtypes, Physiology and Pharmacology: New Ligands and Concepts in Schizophrenia. Front. Pharmacol. 2020, 11. 10.3389/fphar.2020.01003.

(5) Latif, S.; Jahangeer, M.; Maknoon Razia, D.; Ashiq, M.; Ghaffar, A.; Akram, M.; El Allam, A.; Bouyahya, A.; Garipova, L.; Ali Shariati, M.; Thiruvengadam, M.; Azam Ansari, M. Dopamine in Parkinson’s Disease. Clinica Chimica Acta 2021, 522, 114–126. 10.1016/j.cca.2021.08.009.

(6) Seeman, P.; Niznik, H. B. Dopamine Receptors and Transporters in Parkinson’s Disease and Schizophrenia. The FASEB Journal 1990, 4 (10), 2737–2744. 10.1096/fasebj.4.10.2197154.

(7) Foll, B. L.; Gallo, A.; Strat, Y. L.; Lu, L.; Gorwood, P. Genetics of Dopamine Receptors and Drug Addiction: A Comprehensive Review. Behavioural Pharmacology 2009, 20 (1), 1. 10.1097/FBP.0b013e3283242f05.

(8) Jones-Tabah, J.; Mohammad, H.; Paulus, E. G.; Clarke, P. B. S.; Hébert, T. E. The Signaling and Pharmacology of the Dopamine D1 Receptor. Front Cell Neurosci 2022, 15, 806618. 10.3389/fncel.2021.806618.

(9) Szymanski, M. W.; Richards, J. R. Fenoldopam. In StatPearls; StatPearls Publishing: Treasure Island (FL), 2025.

(10) Biglan, K.; Munsie, L.; Svensson, K. A.; Ardayfio, P.; Pugh, M.; Sims, J.; Brys, M. Safety and Efficacy of Mevidalen in Lewy Body Dementia: A Phase 2, Randomized, Placebo-Controlled Trial. Movement Disorders 2022, 37 (3), 513–524. 10.1002/mds.28879.

(11) Bezard, E.; Gray, D.; Kozak, R.; Leoni, M.; Combs, C.; Duvvuri, S. Rationale and Development of Tavapadon, a D1/D5-Selective Partial Dopamine Agonist for the Treatment of Parkinson’s Disease. CNS Neurol Disord Drug Targets 2024, 23 (4), 476–487. 10.2174/1871527322666230331121028.

(12) Gilbert, D. L.; Murphy, T. K.; Jankovic, J.; Budman, C. L.; Black, K. J.; Kurlan, R. M.; Coffman, K. A.; McCracken, J. T.; Juncos, J.; Grant, J. E.; Chipkin, R. E. Ecopipam, a D1 Receptor Antagonist, for Treatment of Tourette Syndrome in Children: A Randomized, Placebo-Controlled Crossover Study. Mov Disord 2018, 33 (8), 1272–1280. 10.1002/mds.27457.

(13) Missale, C.; Nash, S. R.; Robinson, S. W.; Jaber, M.; Caron, M. G. Dopamine Receptors: From Structure to Function. Physiol. Rev. 1998, 78 (1), 189–225. 10.1152/physrev.1998.78.1.189.

(14) Tichy, A.-M.; Gerrard, E. J.; Sexton, P. M.; Janovjak, H. Light-Activated Chimeric GPCRs: Limitations and Opportunities. Current Opinion in Structural Biology 2019, 57, 196–203. 10.1016/j.sbi.2019.05.006.

(15) Whissell, P. D.; Tohyama, S.; Martin, L. J. The Use of DREADDs to Deconstruct Behavior. Front. Genet. 2016, 7. 10.3389/fgene.2016.00070.

(16) Donthamsetti, P.; Winter, N.; Hoagland, A.; Stanley, C.; Visel, M.; Lammel, S.; Trauner, D.; Isacoff, E. Cell Specific Photoswitchable Agonist for Reversible Control of Endogenous Dopamine Receptors. Nat Commun 2021, 12 (1), 4775. 10.1038/s41467-021-25003-w.

(17) Levitz, J.; Broichhagen, J.; Leippe, P.; Konrad, D.; Trauner, D.; Isacoff, E. Y. Dual Optical Control and Mechanistic Insights into Photoswitchable Group II and III Metabotropic Glutamate Receptors. Proc Natl Acad Sci U S A 2017, 114 (17), E3546–E3554. 10.1073/pnas.1619652114.

(18) Broichhagen, J.; Frank, J. A.; Trauner, D. A Roadmap to Success in Photopharmacology. Acc. Chem. Res. 2015, 48 (7), 1947–1960. 10.1021/acs.accounts.5b00129.

(19) Hagen, V.; Kilic, F.; Schaal, J.; Dekowski, B.; Schmidt, R.; Kotzur, N. [8-[Bis(Carboxymethyl)Aminomethyl]-6-Bromo-7-Hydroxycoumarin-4-Yl]Methyl Moieties as Photoremovable Protecting Groups for Compounds with COOH, NH2, OH, and C=O Functions. J. Org. Chem. 2010, 75 (9), 2790–2797. 10.1021/jo100368w.

(20) Lee, T. H.; Gee, K. R.; Ellinwood, E. H.; Seidler, F. J. Combining ‘Caged-Dopamine’ Photolysis with Fast-Scan Cyclic Voltammetry to Assess Dopamine Clearance and Release Autoinhibition in Vitro. J. Neurosci. Methods 1996, 67 (2), 221–231. 10.1016/0165-0270(96)00068-4.

(21) Araya, R.; Andino-Pavlovsky, V.; Yuste, R.; Etchenique, R. Two-Photon Optical Interrogation of Individual Dendritic Spines with Caged Dopamine. ACS Chem. Neurosci. 2013, 4 (8), 1163–1167. 10.1021/cn4000692.

(22) Donthamsetti, P. C.; Broichhagen, J.; Vyklicky, V.; Stanley, C.; Fu, Z.; Visel, M.; Levitz, J. L.; Javitch, J. A.; Trauner, D.; Isacoff, E. Y. Genetically Targeted Optical Control of an Endogenous G Protein-Coupled Receptor. J. Am. Chem. Soc. 2019, 141 (29), 11522–11530. 10.1021/jacs.9b02895.

(23) Donthamsetti, P. C.; Winter, N.; Schönberger, M.; Levitz, J.; Stanley, C.; Javitch, J. A.; Isacoff, E. Y.; Trauner, D. Optical Control of Dopamine Receptors Using a Photoswitchable Tethered Inverse Agonist. J. Am. Chem. Soc. 2017, 139 (51), 18522–18535. 10.1021/jacs.7b07659.

(24) Hetzler, B. E.; Donthamsetti, P.; Peitsinis, Z.; Stanley, C.; Trauner, D.; Isacoff, E. Y. Optical Control of Dopamine D2-like Receptors with Cell-Specific Fast-Relaxing Photoswitches. J. Am. Chem. Soc. 2023, 145 (34), 18778–18788. 10.1021/jacs.3c02735.

(25) Vendrell-Llopis, N.; Read, J.; Boggiano, S.; Hetzler, B.; Peitsinis, Z.; Stanley, C.; Visel, M.; Trauner, D.; Donthamsetti, P.; Carmena, J.; Lammel, S.; Isacoff, E. Y. Dopamine D1 Receptor Activation in the Striatum Is Sufficient to Drive Reinforcement of Anteceding Cortical Patterns. Neuron 2025, 113 (5), 785–794.e9. 10.1016/j.neuron.2024.12.013.

(26) Pendleton, R. G.; Samler, L.; Kaiser, C.; Ridley, P. T. Studies on Renal Dopamine Receptors with a New Agonist. European Journal of Pharmacology 1978, 51 (1), 19–28. 10.1016/0014-2999(78)90057-2.

(27) Setler, P. E.; Sarau, H. M.; Zirkle, C. L.; Saunders, H. L. The Central Effects of a Novel Dopamine Agonist. Eur. J. Pharmacol. 1978, 50 (4), 419–430. 10.1016/0014-2999(78)90148-6.

(28) Iorio, L. C.; Barnett, A.; Leitz, F. H.; Houser, V. P.; Korduba, C. A. SCH 23390, a Potential Benzazepine Antipsychotic with Unique Interactions on Dopaminergic Systems. J Pharmacol Exp Ther 1983, 226 (2), 462–468.

(29) Pettersson, I.; Liljefors, T.; Boegesoe, K. Conformational Analysis and Structure-Activity Relationships of Selective Dopamine D-1 Receptor Agonists and Antagonists of the Benzazepine Series. J. Med. Chem. 1990, 33 (8), 2197–2204. 10.1021/jm00170a025.

(30) Brogden, R. N.; Markham, A. Fenoldopam: A Review of Its Pharmacodynamic and Pharmacokinetic Properties and Intravenous Clinical Potential in the Management of Hypertensive Urgencies and Emergencies. Drugs 1997, 54 (4), 634–650. 10.2165/00003495-199754040-00008.

(31) Kinter, L. B.; Horner, E.; Mann, W. A.; Weinstock, J.; Ruffolo, R. R. Characterization of the Hemodynamic Activities of Fenoldopam and Its Enantiomers in the Dog. Chirality 1990, 2 (4), 219–225. 10.1002/chir.530020405.

(32) Neumeyer, J. L.; Kula, N. S.; Baldessarini, R. J.; Baindur, N. Stereoisomeric Probes for the D1 Dopamine Receptor: Synthesis and Characterization of R-(+) and S-(-) Enantiomers of 3-Allyl-7,8-Dihydroxy-1-Phenyl-2,3,4,5-Tetrahydro-1H-3-Benzazepine and Its 6-Bromo Analog. J. Med. Chem. 1992, 35 (8), 1466–1471. 10.1021/jm00086a016.

(33) Kaiser, C.; Dandridge, P. A.; Garvey, E.; Hahn, R. A.; Sarau, H. M.; Setler, P. E.; Bass, L. S.; Clardy, J. Absolute Stereochemistry and Dopaminergic Activity of Enantiomers of 2,3,4,5-Tetrahydro-7,8-Dihydroxy-1-Phenyl-1H-3-Benzazepine. J. Med. Chem. 1982, 25 (6), 697–703. 10.1021/jm00348a017.

(34) Ohlstein, E. H.; Berkowitz, B. A. SCH 23390 and SK&F 83566 Are Antagonists at Vascular Dopamine and Serotonin Receptors. European Journal of Pharmacology 1985, 108 (2), 205–208. 10.1016/0014-2999(85)90728-9.

(35) Bakthavachalam, V.; Baindur, N.; Madras, B. K.; Neumeyer, J. L. Fluorescent Probes for Dopamine Receptors: Synthesis and Characterization of Fluorescein and 7-Nitrobenz-2-Oxa-1,3-Diazol-4-Yl Conjugates of D-1 and D-2 Receptor Ligands. J Med Chem 1991, 34 (11), 3235–3241. 10.1021/jm00115a012.

(36) Neumeyer, J. L.; Baindur, N.; Bakthavachalam, V.; Yuan, J.; Gao, Y.; Kula, N. S.; Campbell, A.; Baldessarini, R. J. Stereoisomeric, Photoaffinity, Affinity and Fluorescent Probes for Characterization of Dopamine D1 and D2 Receptors. In Pharmacochemistry Library; Angeli, P., Gulini, U., Quaglia, W., Eds.; Trends in Receptor Research; Elsevier, 1992; Vol. 18, pp 175–184. 10.1016/B978-0-444-88931-7.50016-5.

(37) Monsma Jr., F. J.; Barton, A. C.; Chol Kang, H.; Brassard, D. L.; Haugland, R. P.; Sibley, D. R. Characterization of Novel Fluorescent Ligands with High Affinity for D1 and D2 Dopaminergic Receptors. Journal of Neurochemistry 1989, 52 (5), 1641–1644. 10.1111/j.1471-4159.1989.tb09220.x.

(38) Morstein, J.; Awale, M.; Reymond, J.-L.; Trauner, D. Mapping the Azolog Space Enables the Optical Control of New Biological Targets. ACS Cent. Sci. 2019, 5 (4), 607–618. 10.1021/acscentsci.8b00881.

(39) Zhuang, Y.; Xu, P.; Mao, C.; Wang, L.; Krumm, B.; Zhou, X. E.; Huang, S.; Liu, H.; Cheng, X.; Huang, X.-P.; Shen, D.-D.; Xu, T.; Liu, Y.-F.; Wang, Y.; Guo, J.; Jiang, Y.; Jiang, H.; Melcher, K.; Roth, B. L.; Zhang, Y.; Zhang, C.; Xu, H. E. Structural Insights into the Human D1 and D2 Dopamine Receptor Signaling Complexes. Cell 2021, 184 (4), 931–942.e18. 10.1016/j.cell.2021.01.027.

(40) Neumeyer, J. L.; Baindur, N.; Yuan, J.; Booth, G.; Seeman, P.; Niznik, H. B. Development of a High Affinity and Stereoselective Photoaffinity Label for the D-1 Dopamine Receptor: Synthesis and Resolution of 7-[125I]Iodo-8-Hydroxy-3-Methyl-1-(4’-Azidophenyl)-2,3,4,5-Tetrahydro-1H-3-Benzazepine. J. Med. Chem. 1990, 33 (2), 521–526. 10.1021/jm00164a009.

(41) Gautier, A.; Juillerat, A.; Heinis, C.; Corrêa, I. R.; Kindermann, M.; Beaufils, F.; Johnsson, K. An Engineered Protein Tag for Multiprotein Labeling in Living Cells. Chem. Biol. 2008, 15 (2), 128–136. 10.1016/j.chembiol.2008.01.007.

(42) Robertson, G. S.; Robertson, H. A. Synergistic Effects of D1 and D2 Dopamine Agonists on Turning Behaviour in Rats. Brain Research 1986, 384 (2), 387–390. 10.1016/0006-8993(86)91178-9.

(43) Keefe, K. A.; Gerfen, C. R. D1–D2 Dopamine Receptor Synergy in Striatum: Effects of Intrastriatal Infusions of Dopamine Agonists and Antagonists on Immediate Early Gene Expression. Neuroscience 1995, 66 (4), 903–913. 10.1016/0306-4522(95)00024-D.

