## Supplement for "Reversible Antagonism of Dopamine D1 Receptor using a Photoswitchable Remotely Tethered Ligand"

### Supporting Information – P-D1block

#### 1 Supporting Figures

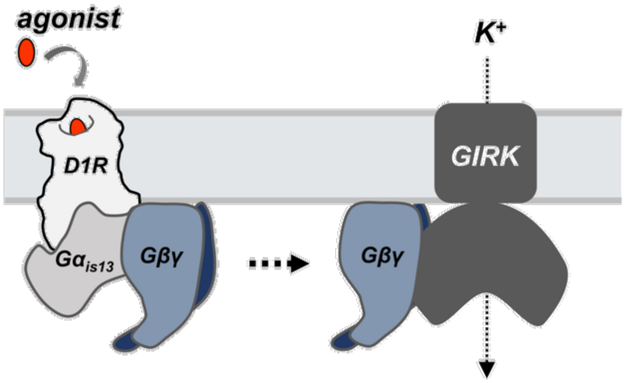


**Supplementary Figure 1. An electrophysiology-based assay for measuring D1R activation.** Schematic representation of a D1R-mediated G protein-coupled inwardly rectifying potassium (GIRK) channel assay in HEK293T cells. Agonist-induced receptor activation results in the recruitment of a heterotrimeric G protein containing the chimeric Gα subunit ($G\alpha_{is13}$) followed by the release of Gβγ, which activates GIRK channels and enhances inward-current.

### 2 Experimental Details

#### 2.1 General Methods, Reagents, Instrumentation

##### 2.1.1 Reagents and Solvents

All reagents and solvents were purchased from commercial sources (Acros Organics, Alfa Aesar, Cayman, Combi-Blocks, Oakwood, Sigma Aldrich, TCI, TRC, etc.) and were used without further purification. Monodisperse PEG-linkers were purchased from PurePEG. Solvents were obtained from Fisher Scientific.

##### 2.1.2 TLC, FCC, LCMS, NMR, FTIR, HPLC

Reactions were monitored by thin layer chromatography (**TLC**) on glass plates precoated with silica gel (0.25 mm, 60 Å pore size, Merck). The plates were visualized by exposure to UV light (254 nm).

Flash silica gel chromatography (**FCC**) was performed on a CombiFlash EZ PrepTM using silica gel ($SiO_2$, particle size 40-63 µm) purchased from SiliCycle.

**NMR** spectra were measured on a Bruker AV-III HD 400 MHz (equipped with a CryoProbeTM) (operating at 400 MHz for $^{1}H$ and 100 MHz for $^{13}C$) or on a Bruker AVIII-600 High Performance Digital NMR Spectrometer with a CPTCI-cryoprobe head (600 MHz for $^{1}H$ and 150 MHz for $^{13}C$). Multiplicities in the following experimental procedures are abbreviated as follows: s = singlet, d = doublet, t = triplet, q = quartet, m = multiplet. $^{1}H$ chemical shifts are expressed in parts per million (ppm, δ scale). The residual protium in the deuterated solvent was used as internal reference (MeOD: δ = 7.26 or $CDCl_3$: δ = 7.26). $^{13}C$ chemical shifts are expressed in ppm (δ scale) and are referenced to the carbon resonance of the NMR solvent (MeOD: δ = 49.00 or $CDCl_3$: δ = 77.16). Structural analysis was conducted with $^{1}H$- and $^{13}C$-NMR spectra using additional 2D spectra (COSY, HMBC, HSQC).

High-Resolution Mass Spectra (**HRMS**) were recorded on an Agilent 6224 Accurate-Mass TOF/LC/MS using an electrospray ionization source (ESI).

**FTIR** were recorded on a ThermoScientific Nicolet-6700 Fourier Transform Infrared Spectroscopy system.

**LCMS** analysis was performed on an LCMS 1260 Infinity II Agilent Technologies system (Windows 10, OpenLabs CDS Chemstation Software, 6120 Quadrupole LC/MS G7111B quaternary pump, G7129A Infinity II vialsampler, G7117C 1260 diode array detector) with an LC Kinetex column 2.6 μm C18 (50 x 3 mm). Runs were performed at a flow-rate of 1 mL/min with a run-time of 5 min, and a solvent gradient of 0-100% MeCN in water, containing 0.1% formic acid.

Preparative **HPLC** was performed on a 1260 Infinity Agilent Technologies system (Windows 10, OpenLabs CDS Chemstation Software, two G1361A pumps, G2260A autosampler (2400 μ L max. injection volume), G1170A column switching valve, G7115A diode array detector, G1364B fraction collector, using a semipreparative column (Phenomenex, Gemini 5 um C18 110 Å, 15- x 10 mm, product #00F-4435-N0) or a preparative column (Phenomenex, Gemini 5 um C18 110 Å, product #00F-4435-U0-AX). Runs were performed at a flowrate of 9 or 80 mL/min (if not specified otherwise), using solvent mixtures of MeCN in $H_2O$, containing 0.1% formic acid.

**Chiral HPLC** was performed on an Agilent 1260 Infinity HPLC on chiral stationary phase (CHIRALCEL OD-H, 4.6 x 250 mm, 24 °C), a flow-rate of 0.8 mL/min using isocratic mobile phase 4% i-PrOH in hexane and under detection at 280 nm.

##### 2.1.2 UV-Vis and Photophysical Characterization

**UV-Vis** spectroscopy was performed on a Varian Cary 60 UV-Visible Spectrometer equipped with an Agilent Technologies PCB 1500 Water Peltier system for temperature control. Samples were measured using disposable Spectrometer/Photometer Ultra-Micro Cuvettes from BrandTech (10 mm light path, 1 mL sample) and irradiation was performed with a Cairn Research Optoscan Monochromator with Optosource High Intensity Arc Lamp equipped with a 75 W UXL-S50A lamp from USHIO Inc. Japan, set to 15 nm full width at half maximum. The Monochromator was controlled using a MATLAB program written by Christopher Arp. UV-Vis data were analyzed and plotted using GRAPHPAD Prism. Irradiation was achieved from the top of a cuvette through a fiber-optic cable. Reversible switching was performed by diluting samples to 20 µM and irradiating with 415/600 nm for 90 seconds each. Absorbance (reported as Abs, in arbitrary units) was measured at 420 nm over time. Thermal Relaxation was measured after 1 min irradiation with 415 nm at 10 or 20 µM, and subsequently absorption increase was detected at 420 nm. The relaxation half-live of the *cis*-isomer was determined by curve fitting, using exponential one-phase decay in GraphPad Prism. Photo-stationary states were determined by pre-irradiation of samples (20 uM, DMSO) for 10 min at the respective wavelengths. Samples for LCMS separation were prepared under red-light conditions in amber vials and immediately subjected to LCMS. The relative ratios of (*Z*)- and (*E*)-isomers ($t_R$ = 2.662 min and $t_R$ = 2.748 min) were determined by detection at the isosbestic point at the respective elution time solvent mixtures (305 nm).

##### 2.1.3 Molecular Biology and Heterologous Expression

All constructs were cloned into mammalian expression vectors and are available from the authors upon reasonable request. For the receptor mediated-GIRK activation assay, HEK293T cells were seeded onto 18 mm coverslips and transiently transfected overnight with Lipofectamine 2000 and the following constructs: D1R (0.525 µg), the membrane-anchor protein (the M protein; 0.7 µg), $G\alpha_{is13}$ (0.35 µg), GIRK1(F137S) (0.7 µg), and tdTomato (0.2 µg). Transfected cells were used for electrophysiology experiments.

##### 2.1.4 Electrophysiology

HEK293T cells were sparsely seeded and maintained in DMEM (Invitrogen) with 10% fetal bovine serum on poly-L-lysine-coated coverslips at 37°C and 5% $CO_2$. HEK293T cells were voltage clamped in whole-cell configuration 16-48 hours after transfection. For GIRK experiments, the extracellular solution contained 120 mM KCl, 25 mM NaCl, 10 mM HEPES, 2 mM CaCl2, and 1 mM MgCl2, pH 7.4. Glass pipettes with a resistance of 3-7 MΩ were filled with intracellular solution containing 120 mM Gluconic acid δ-lactone, 15 mM CsCl, 10 mM BAPTA, 10 mM HEPES, 1 mM $CaCl_2$, 3 $MgCl_2$, 3 mM MgATP, pH 7.2. Cells were voltage clamped to -80 mV using an Axopatch 200A (Molecular Devices) amplifier.

To conjugate P-$D1_{block}$ variants to the M protein, cells were incubated with 1 µM compound for 60 minutes in the dark at 37°C in standard extracellular buffers. For all experiments, compounds were applied using a gravity-driven perfusion system and illumination was applied to the entire field of view using a DG4 (Sutter) through a 20x objective (0.5 $mW/mm^2$ at 370 nm and 1.6 $mW/mm^2$ at 460 nm). pClamp software was used for both data acquisition and control of illumination.

The selection criteria for electrophysiological experiments are that a cell (i) expresses the fluorescent protein transfection marker, and (ii) responds to agonist, indicating the presence of either receptor and GIRK. Cells were not excluded unless the recording was of poor quality (e.g., unstable baseline).

#### 2.2 Chemistry

##### 2.2.1 2-(4-nitrophenyl)oxirane 2

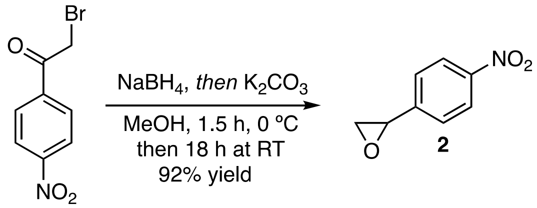

1

2-(4-nitrophenyl)oxirane (**2**) was prepared according to the procedure from Vollrath, Benedikt et al from PCT Int. Appl., 201044535, 14 Apr 2011.In a 250 mL flame-dried rbf under magnetic stirring, 2-bromo-1-(4-nitrophenyl)ethan-1-one (**1**) (5.0 g, 20.5 mmol, 1.0 eq) was dissolved in anhydrous MeOH (50 mL, 0.4M) and cooled to 0 °C. $NaBH_4$ (853 mg, 22.5 mmol, 1.10 eq) was added to the reaction mixture at 0 °C in small portions under gas evolution, and the resulting mixture was stirred for 1.5 h until full conversion was determined by TLC. $K_2CO_3$ (2.8 g, 20.5 mmol, 1.0 eq) was then added at 0 °C and the resulting mixture stirred at room temperature overnight. The reaction mixture was diluted with brine (40 mL) and extracted with diethyl ether (3x 100 mL). The combined organic phases were dried over $Na_2SO_4$ and solvent removed under reduced pressure to yield the epoxide **2** as orange solid in 92% (3.12 g, 18.9 mmol).

**R$_f$** = 0.61 (EtOAc/hexanes, 1:1; UV).

**LCMS** (5-100% MeCN in $H_2O$ with 0.1% formic acid over 5 min) $t_R$ = 3.057 min, 230 nm detection.

**LRMS** (ESI): calc. for $C_8H_7NO_3^+$ $[M+H]^+$: 166.0; found 166.1.

**$^1$H NMR** (400 MHz, $CDCl_3$): δ = 8.2 – 8.2 (m, 2H), 7.5 – 7.4 (m, 2H), 4.0 (dd, J = 4.1, 2.5 Hz, 1H), 3.2 (dd, J = 5.5, 4.1 Hz, 1H), 2.8 (dd, J = 5.5, 2.4 Hz, 1H).

**$^{13}$C NMR** (101 MHz, $CDCl_3$): δ = 148.0, 145.4, 126.4, 124.0, 51.8, 51.6.

**IR** (neat) 1606 (w), 1519 (s), 1476 (w), 1344 (s), 1316 (w), 1203 (w), 1107 (w), 988 (m), 876 (m), 847 (s), 749 (m) cm$^{-1}$.

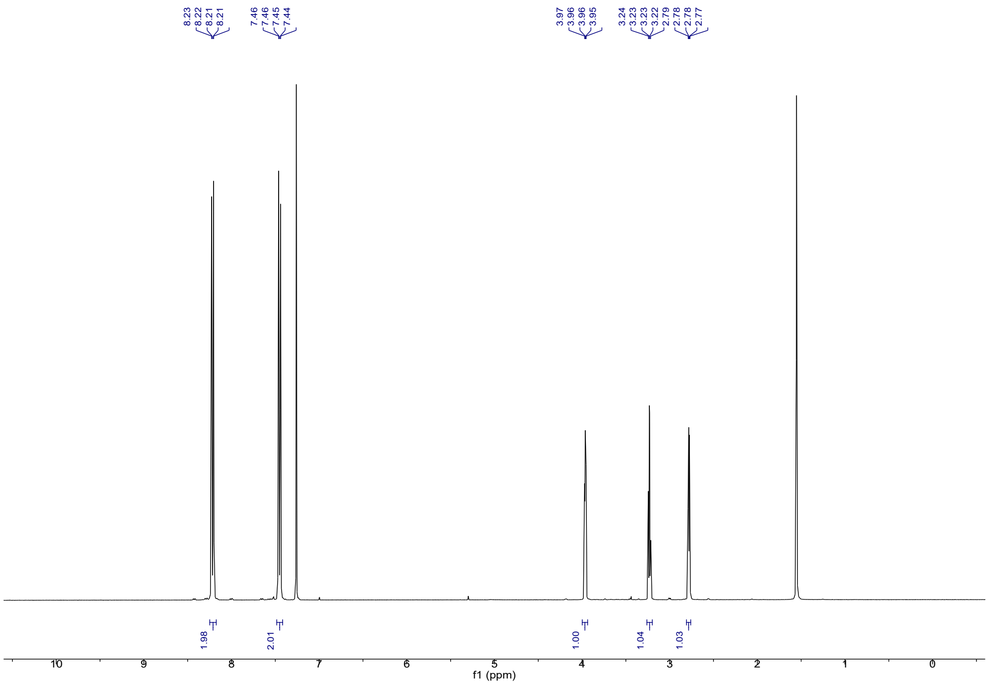

8.23
8.22
8.21
8.21
7.46
7.46
7.45
7.44
3.97
3.96
3.96
3.95
3.24
3.23
3.23
3.22
2.79
2.78
2.78
2.77
1.98
2.01
1.00
1.04
1.03
10
9
8
7
6
5
4
3
2
1
0
f1 (ppm)


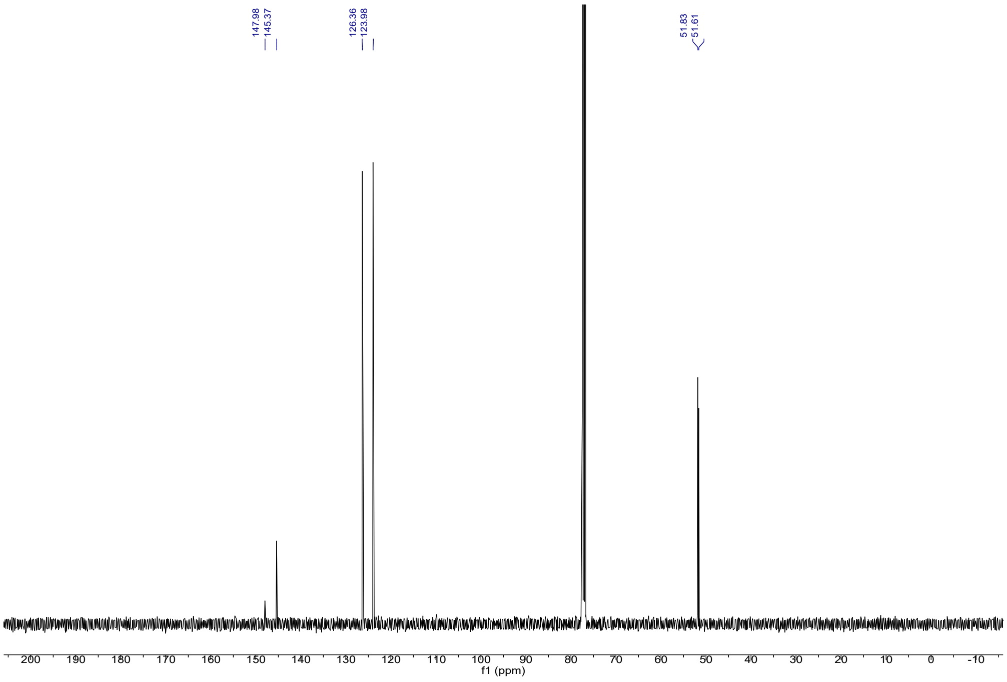

147.98
145.37
126.36
123.98
51.83
51.61
200
190
180
170
160
150
140
130
120
110
100
90
80
70
60
50
40
30
20
10
0
-10
f1 (ppm)

##### 2.2.2 Amino Alcohol 4

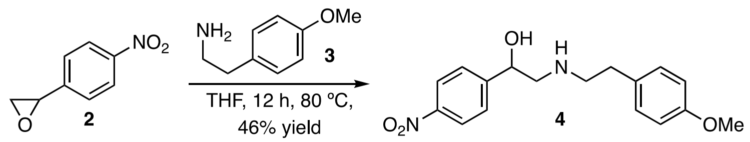


Amino alcohol **4** was prepared according to the procedure from *J. Med. Chem.* **1990**, *33*, 521–526. In a 250 mL flame-dried rbf under magnetic stirring, **2** (2.56 g, 15.5 mmol, 1.0 eq) was dissolved in anhydrous THF (25 mL, 0.6M). 2-(4-methoxyphenyl)ethan-1-amine **3** (2.56 g, 16.9 mmol, 1.1 eq) was added to the reaction flask, and the resulting mixture was stirred with a teflon sleeve and a reflux condenser in a metal heating block at reflux (80 °C heating block temperature) for 16 h. The red reaction solution was then cooled to ambient temperature and the solvent was then removed under reduced pressure. The crude red oil was dissolved in 20 mL of $Et_2O$ and cooled to 0 °C for 30 min, and a white solid formed. The precipitate was filtered off, to yield the amino alcohol **4** as an off-white solid (2.27 g, 7.18 mmol, 46%).

$\mathbf{R}_f$ = 0.36 (5% MeOH in DCM; UV).

**LCMS** (5-100% MeCN in $H_2O$ with 0.1% formic acid over 5 min) $t_R$ = 2.647 min, 230 nm detection.

**LRMS** (ESI): calc. for $C_{17}H_{20}N_2O_4^+$ $[M+H]^+$: 317.1; found 317.1.

**HRMS** (ESI): calc. for $C_{17}H_{20}N_2O_4^+$ $[M+H]^+$: 317.1496; found 317.1498.

**$^1$H NMR** (400 MHz, $CDCl_3$) δ 8.2 – 8.1 (m, 2H), 7.5 (d, *J* = 8.7 Hz, 2H), 7.2 – 7.1 (m, 2H), 6.8 (d, *J* = 8.6 Hz, 2H), 4.9 (dd, *J* = 9.5, 3.4 Hz, 1H), 3.8 (s, 3H), 3.1 – 2.9 (m, 3H), 2.9 – 2.8 (m, 2H), 2.7 (dd, *J* = 12.3, 9.4 Hz, 1H).

**$^{13}$C NMR** (101 MHz, $CDCl_3$): δ = 158.4, 149.2, 147.5, 130.4, 129.6, 126.5, 123.7, 114.1, 69.9, 56.1, 55.3, 50.4, 34.4.

**IR** (neat) 3302 (w), 2896 (w), 2835 (w), 1603 (w), 1509 (s), 1423 (m), 1352 (S), 1240 (s), 1201 (w), 1176 (m), 1107 (s), 1075 (m), 1035 (m), 1014 (w), 986 (w), 906 (m), 884 (m), 856 (m), 837 (m), 818 (s), 753 (m) $cm^{-1}$.

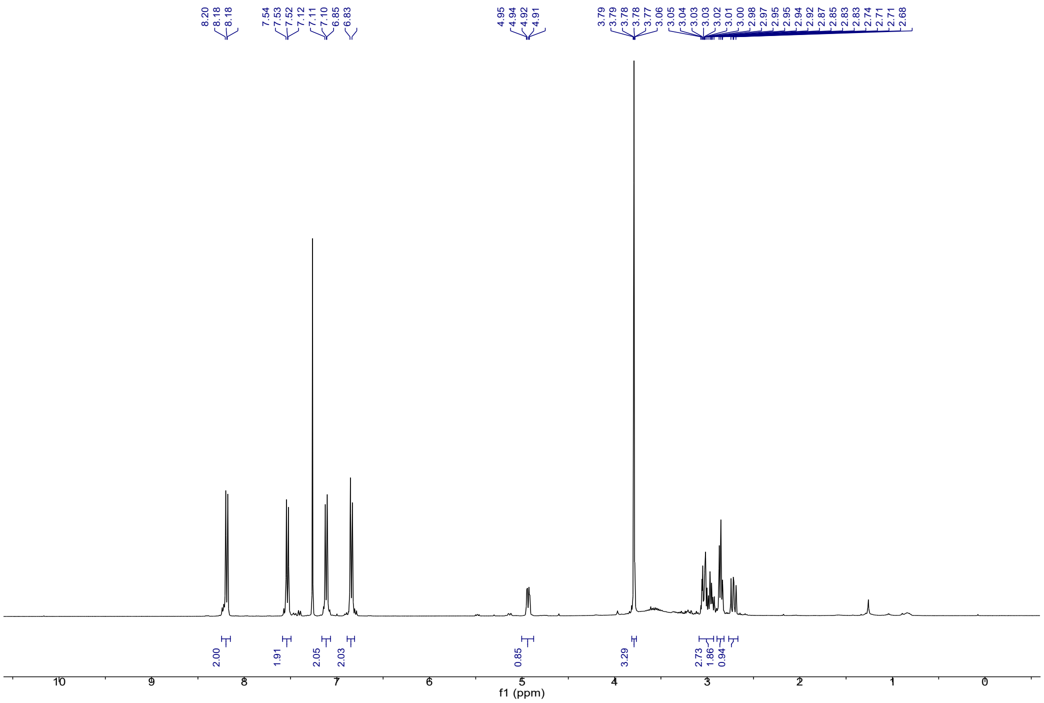
8.20
8.18
8.18
7.54
7.53
7.52
7.12
7.11
7.10
6.85
6.83
4.95
4.94
4.92
4.91
3.79
3.79
3.78
3.78
3.77
3.06
3.05
3.04
3.03
3.03
3.02
3.01
3.00
2.98
2.97
2.95
2.95
2.94
2.92
2.92
2.87
2.85
2.83
2.83
2.74
2.71
2.71
2.68
2.00
1.91
2.05
2.03
0.85
3.29
2.73
1.86
0.94
f1 (ppm)

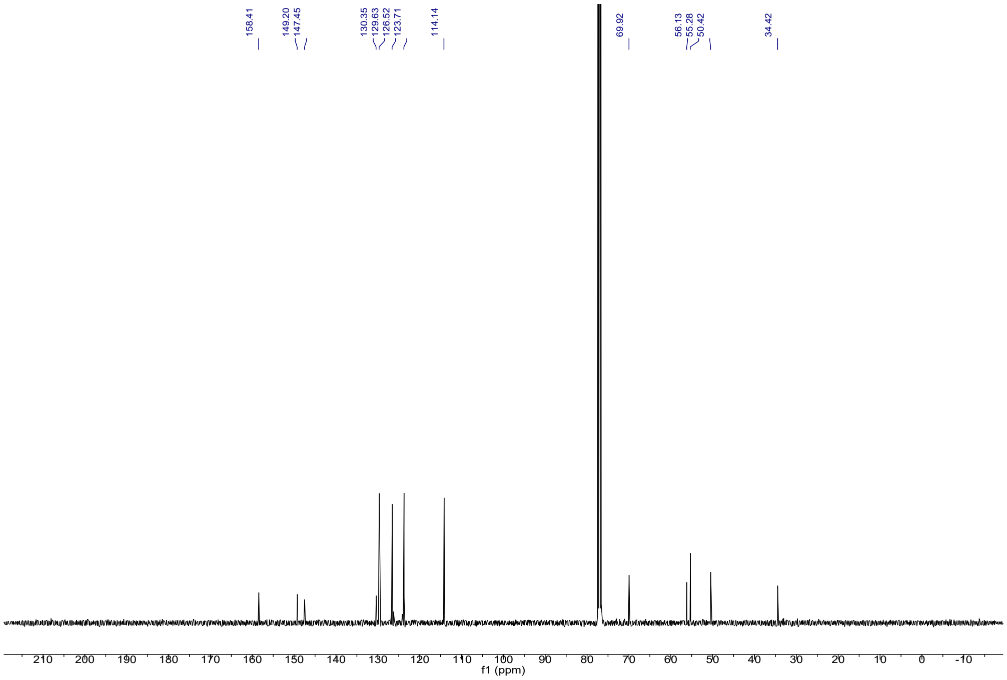
158.41
149.20
147.45
130.35
129.63
126.52
123.71
114.14
69.92
56.13
55.28
50.42
34.42
f1 (ppm)

##### 2.2.3 Nitro Tetrahydro-1H-3-Benzazepine 5

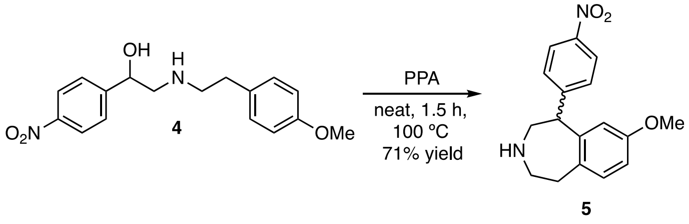


Tetrahydro-1H-3-benzazepine **5** was prepared according to the procedure from *J. Med. Chem.* **1990**, *33*, 521–526. In a 50 mL rbf under magnetic stirring, **4** (1.80 g, 5.69 mmol, 1.0 eq) was combined with polyphosphoric acid (115%, 7.9 mL, 0.7M), heated to 100 °C in an oil bath, and stirred for 2 h. The dark brown reaction mixture was poured while hot into an Erlenmeyer flask with ice-water (10 mL) to form a brown mixture, and then $NH_4OH$ (20 mL) was added to bring the mixture pH above 9. At pH > 9, the product turned red and was extracted with DCM (2x 100 mL), and the combined organic layers dried over $Na_2SO_4$ and concentrated under reduced pressure. The product **5** was recovered as a red solid (1.20 g, 4.02 mmol, 71%) and used without further purification. An analytical sample was purified on normal phase semi prep (0-5% MeOH in DCM over 20 min, 19 mL/min, 20 mm x 250 mm x 5 uM) to yield a pure sample of **5** for characterization.

**R*f*** = 0.28 (5% MeOH in DCM; UV).

**LCMS** (5-100% MeCN in $H_2O$ with 0.1% formic acid over 5 min) $t_R$ = 2.510 min, 360 nm detection.

**LRMS** (ESI): calc. for $C_{17}H_{18}N_2O_3^+$ $[M+H]^+$: 299.1; found 299.1.

**HRMS** (ESI): calc. for $C_{17}H_{18}N_2O_3^+$ $[M+H]^+$: 299.1390; found 299.1395.

**$^1$H NMR** (400 MHz, $CDCl_3$) δ 8.2 (d, *J* = 8.4 Hz, 2H), 7.3 (d, *J* = 8.4 Hz, 2H), 7.1 (d, *J* = 8.3 Hz, 1H), 6.7 (dd, *J* = 8.3, 2.7 Hz, 1H), 6.4 (dd, *J* = 4.8, 2.7 Hz, 1H), 4.4 (d, *J* = 6.8 Hz, 1H), 3.7 – 3.6 (m, 4H), 3.5 – 3.4 (m, 1H), 3.1 – 2.7 (m, 4H).

**$^{13}$C NMR** (101 MHz, $CDCl_3$) δ 158.4, 149.4, 146.7, 143.1, 143.1, 133.2, 131.7, 129.3, 124.0, 116.4, 116.4, 111.2, 55.3, 52.5, 52.4, 48.2, 37.9.

**IR** (neat)) ν =2924 (w), 2835 (w), 1607 (m), 1518 (s), 1462 (w), 1347 (s), 1269 (w), 1108 (w), 1049 (w), 856 (w) $cm^{-1}$.

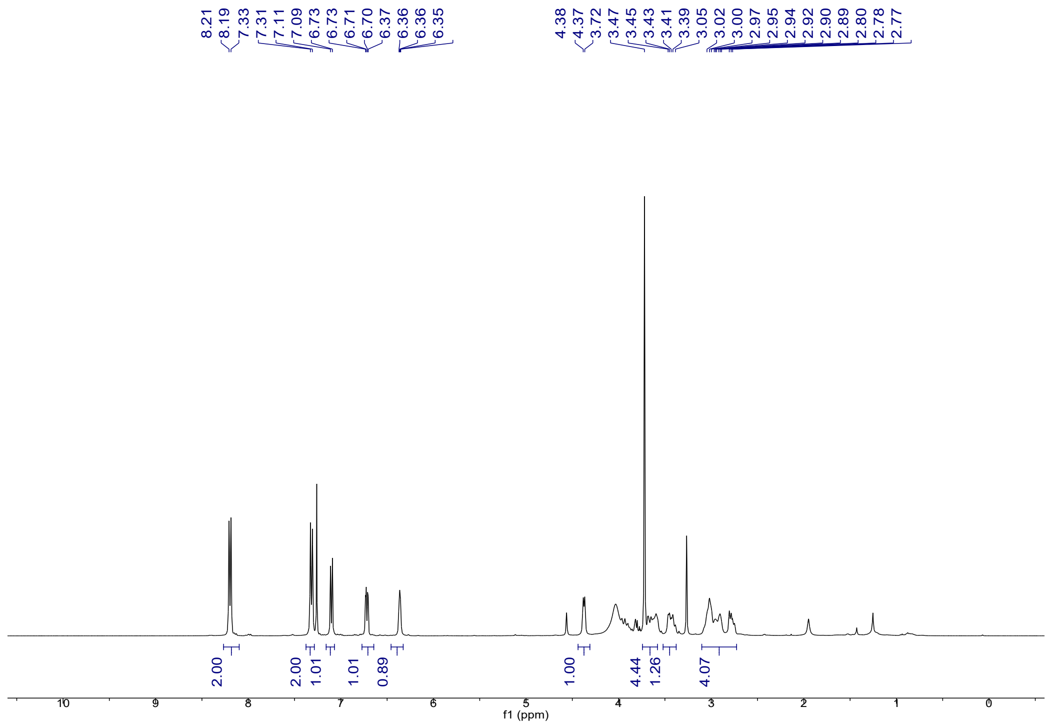
8.21
8.19
7.33
7.31
7.11
7.09
6.73
6.73
6.71
6.70
6.37
6.36
6.36
6.35
4.38
4.37
3.72
3.47
3.45
3.43
3.41
3.39
3.05
3.02
3.00
2.97
2.95
2.94
2.92
2.90
2.89
2.80
2.78
2.77
2.00
2.00
1.01
1.01
0.89
1.00
4.44
1.26
4.07
f1 (ppm)

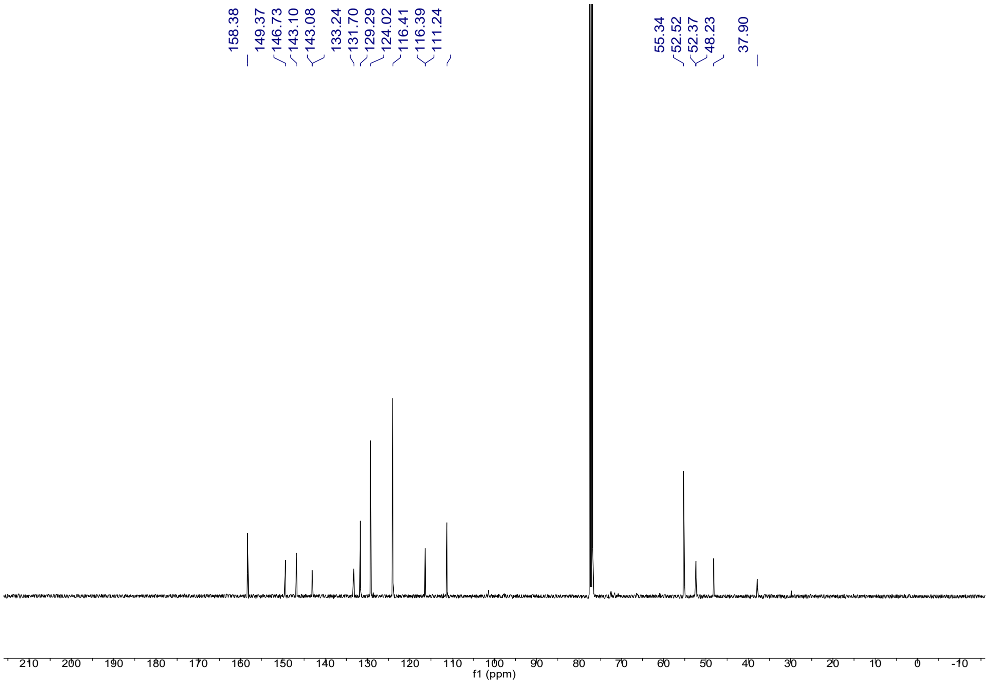
158.38
149.37
146.73
143.10
143.08
133.24
131.70
129.29
124.02
116.41
116.39
111.24
55.34
52.52
52.37
48.23
37.90
f1 (ppm)

##### 2.2.4 N-Methyl Tetrahydro-1H-3-Benzazepine 6

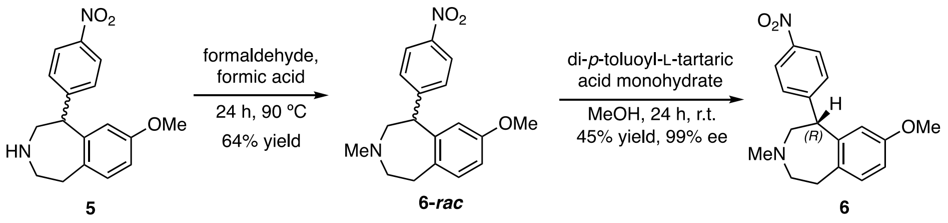


**6** was prepared according to the procedure from *J. Med. Chem.* **1990**, *33*, 521–526. In a 50 mL rbf, **5** (2.60 g, 8.72 mmol, 1.00 eq) was dissolved in formic acid (44 mL) and connected to a nitrogen line via a needle and rubber septum. Then, under stirring, formaldehyde (37% solution in water, stabilized with MeOH; 6.48 mL, 79.8 mmol, 9.16 eq) was added dropwise. The rubber septum was replaced with a Teflon sleeve and a reflux condenser, connected to a nitrogen line. The reaction mixture was heated to 90 °C in an oil-bath for 4 h. The reaction mixture was cooled to room temperature and added dropwise to a 500 mL Erlenmeyer flask containing ice-water (100 mL) and an ammonia solution (aq., 2M, 100 mL) under strong formation of fumes. The white precipitate forming dissolves upon addition of DCM (50 mL). The aq. phase was separated and extracted three times with DCM (3 x 50 mL). The combined organic phases were dried over $Na_2SO_4$, then concentrated *in vacuo*. The racemic *N*-methylated product **6-*rac*** was obtained as a light orange oil in 64% yield (1.75 g, 5.61 mmol).

**Chiral resolution:**

The racemic product **6-*rac*** (2.18 g , 6.98 mmol, 1.00 eq) was dissolved in MeOH (10 mL) in a 100 mL round bottom flask and di-p-toluoyl-L-tartaric acid monohydrate (2.84 g, 7.02 mmol, 1.00 eq) was added. The mixture was warmed in a metal heat block with a reflux condenser under magnetic stirring to 50 °C, until all solid dissolved. The mixture was let stand at ambient temperature for 24 h, until white solid formed, which was harvested over a filter funnel under reduced pressure and washed with little amounts of $Et_2O$. The desired salt was obtained (2.60 g) and recrystallized from MeOH. The obtained salt after recrystallization (1.51 g) was dissolved in water (30 mL) and $NaHCO_3$ (sat., aq., 70 mL). The aqueous phase was extracted three times with EtOAc (50 mL), and the combined, yellow organic phases were washed with brine (20 mL) and dried over $Na_2SO_4$. The organic phase was filtered and concentrated under reduced pressure to yield a red oil, that was subjected to the same procedure. After two rounds of crystallization, the *R*-enantiomer **6** was obtained as a red oil in 30% yield (0.66 g, 2.1 mmol). The combined filtrates containing the undesired *S*-enantiomer were treated identically by free-basing with sodium bicarbonate, to yield 70% of the product enriched in (*S*)-isomer (1.52 g, 4.86 mmol).

Enantiomeric excess of the obtained desired R-(+)-enantiomer was determined by HPLC analysis on chiral stationary phase (CHIRALCEL OD-H, 4.6 x 250 mm, 24 °C, 0.8 mL/min, 4% i-PrOH in hexane, detection at 280 nm) to be 99.5% by comparison with a racemic sample; $t_R$ (*R*-(+)-enantiomer) = 11.702 min, $t_R$ (*S*-(–)-enantiomer) = 14.410 min. The obtained (*S*)-enantiomer in the filtrate had low enantiomeric purity and was treated in the same fashion with di-p-toluoyl-D-tartaric acid monohydrate to yield the pure (*S*)-enantiomer.

**$R_f$** = 0.21 (5% MeOH in DCM; UV).

**LCMS** (5-100% MeCN in $H_2O$ with 0.1% formic acid over 5 min) $t_R$ = 2.605 min, 360 nm detection.

**LRMS** (ESI): calc. for $C_{18}H_{21}N_2O_3^+$ $[M+H]^+$: 313.2; found 313.1.

**HRMS** (APCI): calc. for $C_{18}H_{21}N_2O_3^+$ $[M+H]^+$: 313.1547; found 313.1555.

**$^1$H NMR** (400 MHz, $CDCl_3$) δ 8.2 (d, *J* = 8.7 Hz, 2H), 7.4 – 7.3 (m, 2H), 7.1 (d, *J* = 8.2 Hz, 1H), 6.7 (dd, *J* = 8.2, 2.7 Hz, 1H), 6.3 (d, *J* = 2.7 Hz, 1H), 4.3 (d, *J* = 7.4 Hz, 1H), 3.7 (s, 3H), 3.2 – 3.1 (m, 1H), 3.0 – 2.8 (m, 2H), 2.8 – 2.6 (m, 2H), 2.6 – 2.5 (m, 1H), 2.4 (s, 3H).

**$^{13}$C NMR** (101 MHz, $CDCl_3$): δ = 158.2, 150.5, 146.4, 143.7, 133.5, 131.0, 129.2, 123.7, 115.6, 110.9, 61.8, 57.7, 55.2, 50.4, 48.0, 35.4.

**For (*R*)-enantiomer: $[\alpha]_D$** = + 45.5 (c = 0.5 in MeOH).

**For (*S*)-enantiomer: $[\alpha]_D$** = – 43.4 (c = 0.5 in MeOH).

**IR** (neat) 2937 (w), 2837 (w), 2797 (w), 1652 (w), 1604 (m), 1580 (w), 1517 (s), 1462 (m), 1384 (w), 1346 (s), 1265 (w), 1159 (w), 1108 (m), 1040 (w), 1014 (w), 856 (m), 842 (w), 815 (w), 740 (w) $cm^{-1}$.

**Chiral HPLC of (*R*)-enantiomer:**

**Racemic sample**

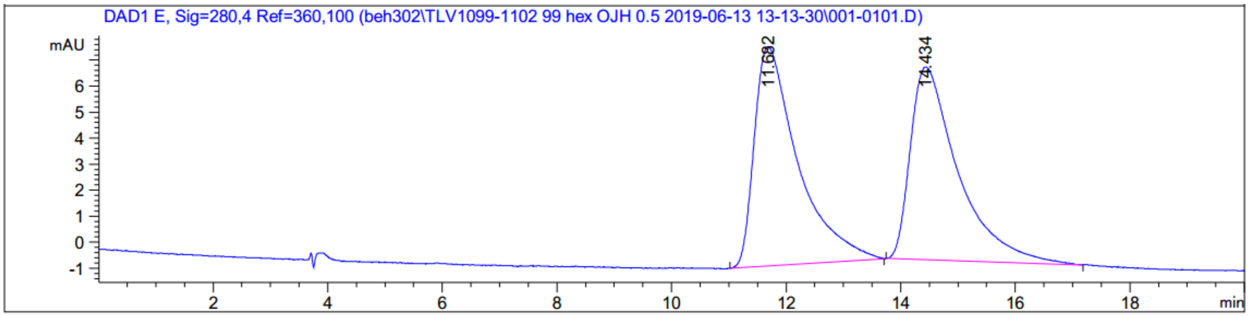


| # | [min] | | [min] | [mAU*s] | [mAU] | % |
|---|---|---|---|---|---|---|
| 1 | 11.682 | BB | 0.7307 | 444.10196 | 8.42872 | 50.9588 |
| 2 | 14.434 | BB | 0.7833 | 427.39038 | 7.40601 | 49.0412 |

**Obtained (*R*)- product after chiral resolution**

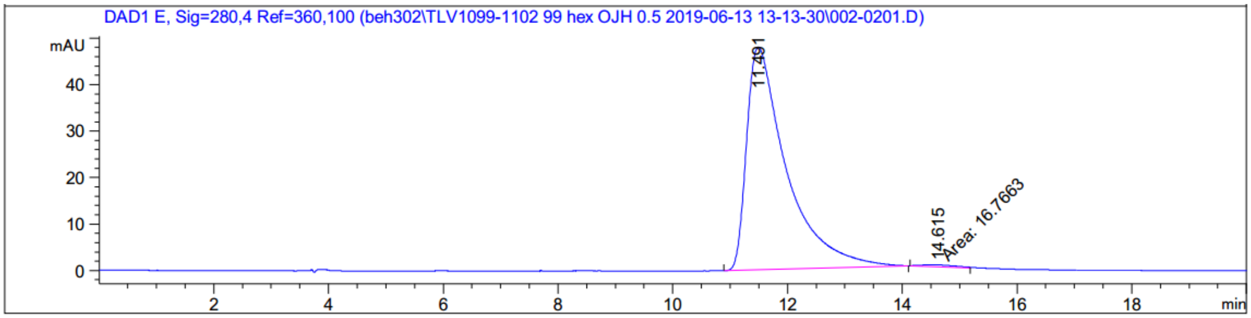


| # | [min] | | [min] | [mAU*s] | [mAU] | % |
|---|---|---|---|---|---|---|
| 1 | 11.491 | BB | 0.6878 | 2307.73657 | 47.92437 | 99.2787 |
| 2 | 14.615 | MM | 0.6817 | 16.76630 | 4.09943e-1 | 0.7213 |

**Co-injection**

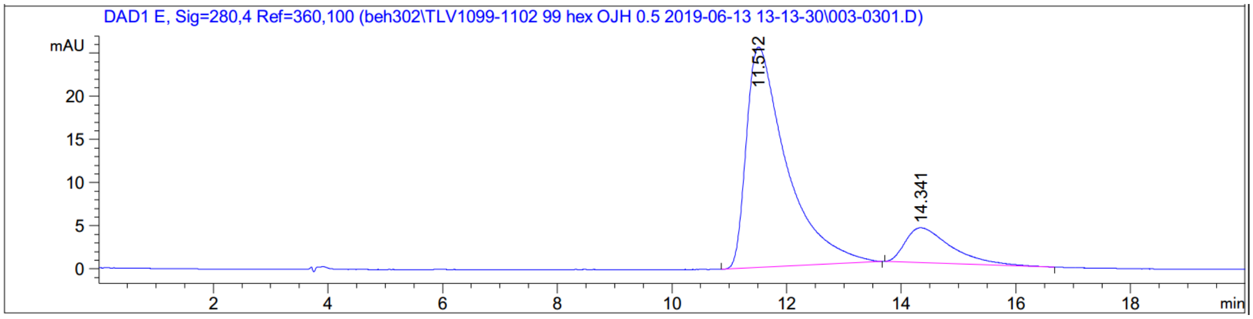


| # | [min] | | [min] | [mAU*s] | [mAU] | % |
|---|---|---|---|---|---|---|
| 1 | 11.512 | BB | 0.6971 | 1246.36560 | 25.55072 | 84.6651 |
| 2 | 14.341 | BB | 0.6571 | 225.74692 | 4.04493 | 15.3349 |

**UV Vis spectra of both enantiomers overlayed**

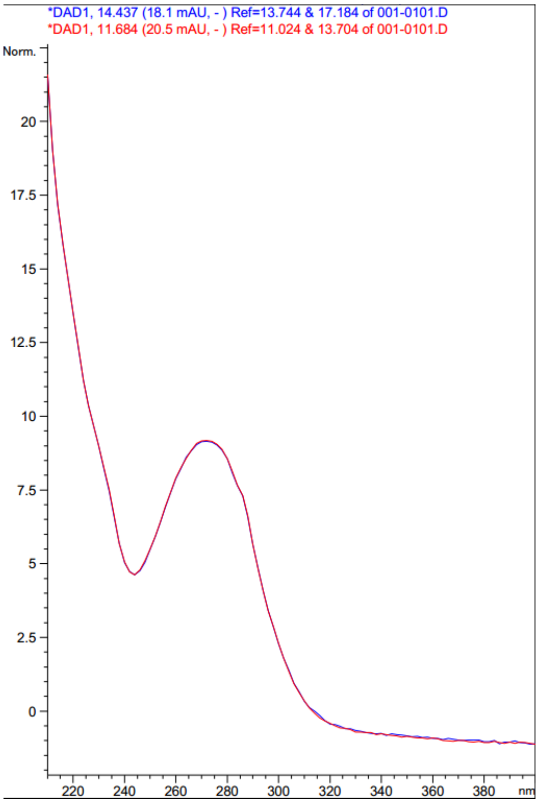

#### [1]H NMR

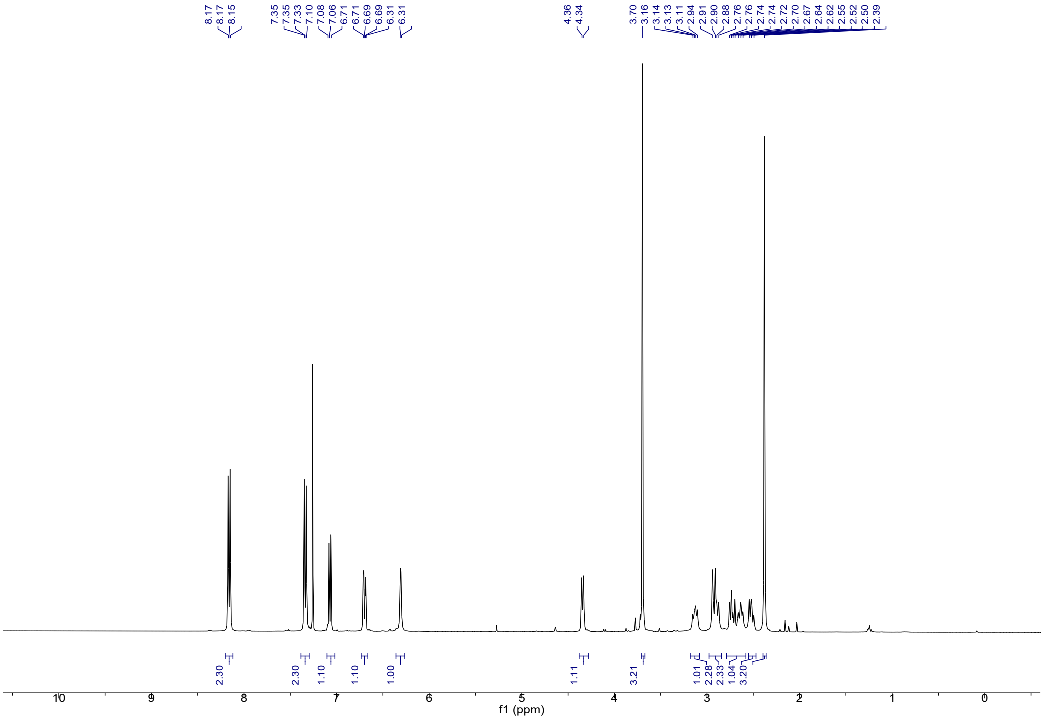


#### [13]C NMR

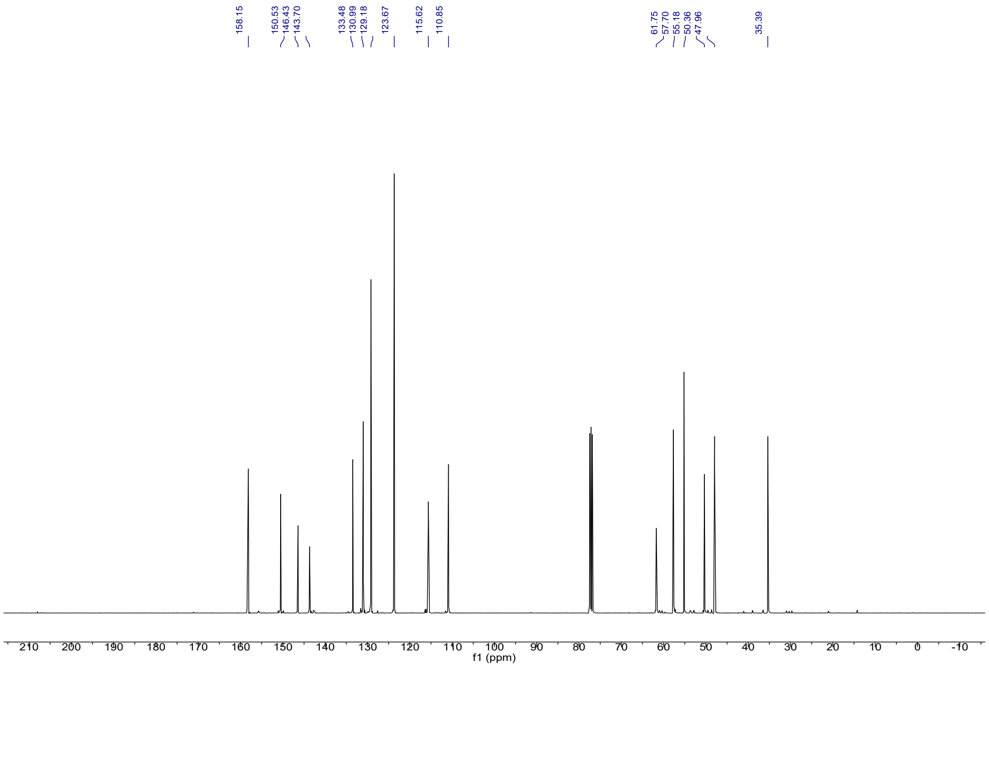

##### 2.2.5 *R*-Bromo Nitro Tetrahydro-1H-3-Benzazepine 7

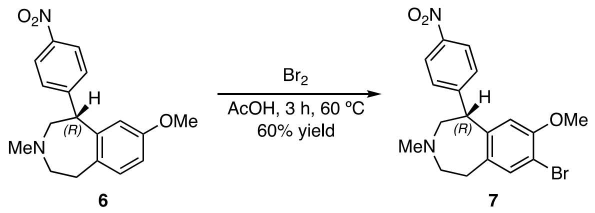


Aryl bromide **7** was prepared according to the procedure from *J. Med. Chem.* **1990**, *33*, 521–526. In a 100 mL three-neck round bottom flask, **6** (0.500 g, 1.60 mmol, 1.00 eq) was dissolved in AcOH (35 mL) and warmed to 60 °C with a reflux condenser and a 250 mL dropping funnel, attached to a Schlenck line under $N_2$ pressure. Bromine (0.460 g, 3.20 mmol, in 10 mL AcOH) was transferred to the dropping funnel and added dropwise at 60 °C. The red solution formed an abundant white precipitate upon addition, which re-dissolved under stirring. After 3 h, the solution was cooled to ambient temperature, poured onto ice-water (50 mL) and treated with ammonium hydroxide (10% aq., 50 mL) and the mixture was extracted with DCM (30 mL x 3). The combined yellow organic phases were washed with brine (20 mL) and dried over $Na_2SO_4$. The solution was filtered and concentrated under reduced pressure, to yield the crude product as an off-yellow foam. The crude product was subjected to flash column chromatography ($SiO_2$, 24g, 0-100% EtOAc in DCM) and the purified bromide **7** was collected as a white foam in 60% yield (377 mg, 0.964 mmol).

**$R_f$** = 0.59 (5% MeOH in DCM; UV).

**LCMS** (5-100% MeCN in $H_2O$ with 0.1% formic acid over 5 min) $t_R$ = 3.011 min, 254 nm detection.

**LRMS** (ESI): calc. for $C_{18}H_{20}BrN_2O_4^+$ $[M+H]^+$: 391.1; found 391.0.

**HRMS** (APCI): calc. for $C_{18}H_{20}BrN_2O_4^+$ $[M+H]^+$: 391.0652; found 391.0654.

**$^1$H NMR** (400 MHz, $CDCl_3$) δ 8.2 – 8.1 (m, 2H), 7.4 – 7.3 (m, 3H), 6.3 (s, 1H), 4.4 (d, *J* = 7.3 Hz, 1H), 3.7 (s, 3H), 3.2 – 3.1 (m, 1H), 2.9 – 2.8 (m, 2H), 2.8 – 2.7 (m, 1H), 2.7 – 2.5 (m, 2H), 2.4 (s, 3H).

$^{13}$**C NMR** (101 MHz, $CDCl_3$) δ 154.4, 149.9, 146.6, 142.8, 134.9, 134.7, 129.1, 123.8, 113.3, 109.6, 61.4, 57.3, 56.3, 50.3, 47.9, 34.9.

**IR** (neat) 3399 (b), 2929 (m), 2854 (w), 1598 (m), 1519 (s), 1495 (m), 1463 (w), 1383 (w), 1346 (s), 1271 (w), 1253 (w), 1110 (w), 1053 (w), 1015 (w), 856 (m) $cm^{-1}$.

**For (*R*)-enantiomer: $[\alpha]_D$** = + 17.2 (c = 0.5 in MeOH).

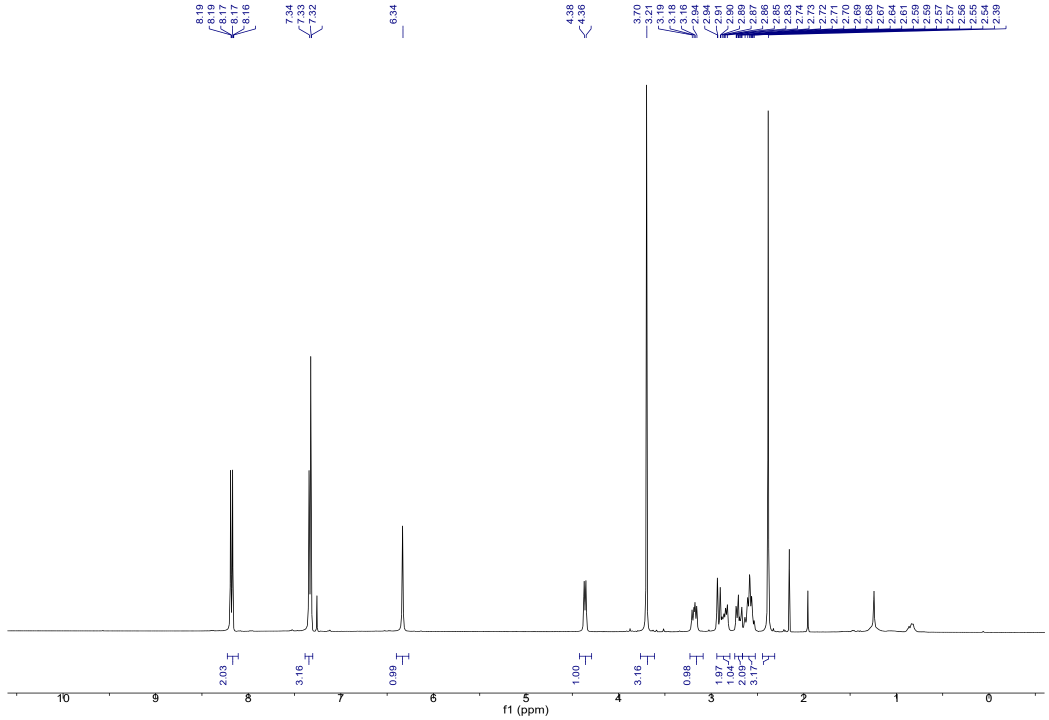

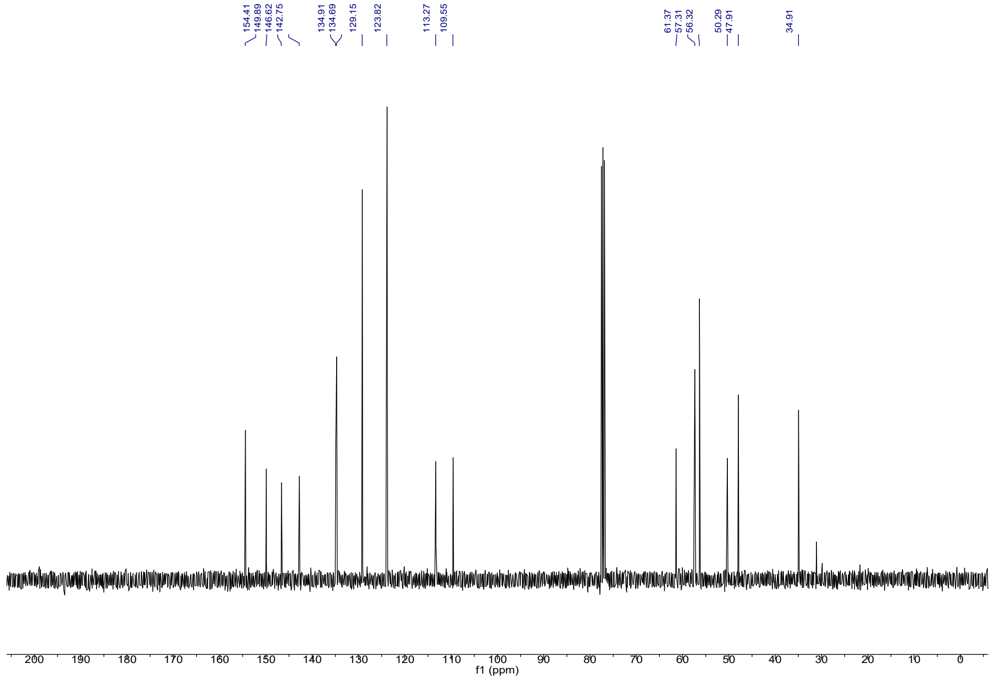

##### 2.2.6 Aniline 8

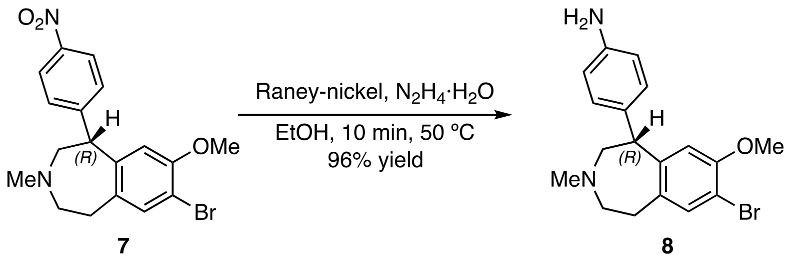

Aniline **8** was prepared according to the procedure from *J. Med. Chem.* **1990**, *33*, 521–526. In a 25 mL round bottom flask, **7** (0.110 g, 0.284 mmol, 1.00 eq) was dissolved in EtOH (5.6 mL), and hydrazine monohydrate (58 µL, 1.19 mmol, 4.20 eq) were added. The reaction mixture was warmed to 50 °C in an oil bath, and Raney Nickel was added as a suspension in water dropwise, until bubbling ceased after about 10 min. The reaction mixture was filtered over celite to remove the Nickel and the crude mixture was concentrated under reduced pressure. The crude yellow oil was subjected to flash column chromatography ($SiO_2$, 4g, 0-5% MeOH in DCM) to yield aniline **8** as light-yellow oil in 96% (98 mg, 0.27 mmol).

$\mathbf{R}_{\boldsymbol{f}}$ = 0.16 (5% MeOH in DCM; UV).

**LCMS** (5-100% MeCN in $H_2O$ with 0.1% formic acid over 5 min) $t_R$ = 1.978 min, 254 nm detection.

**LRMS** (ESI): calc. for $C_{18}H_{21}BrN_2O$ $[M+H]^+$: 361.1; found 361.0.

**HRMS** (APCI): calc for $C_{18}H_{21}BrN_2O$ $[M+H]^+$: 361.0910; found 361.0905.

$^{1}$**H NMR** (400 MHz, $CDCl_3$) δ 7.3 – 7.3 (m, 1H), 7.0 – 6.9 (m, 2H), 6.7 – 6.6 (m, 2H), 6.3 (s, 1H), 4.2 (d, *J* = 8.5 Hz, 1H), 3.6 (s, 3H), 3.1 – 2.9 (m, 2H), 2.9 – 2.7 (m, 3H), 2.4 – 2.3 (m, 4H).

$^{13}$**C NMR** (101 MHz, $CDCl_3$): δ = 154.1, 145.8, 145.0, 134.9, 133.8, 132.7, 129.2, 115.5, 113.0, 108.4, 63.1, 57.3, 56.2, 49.2, 47.8, 35.2.

**For (*R*)-enantiomer: $[\alpha]_D$** = + 28.4 (c = 0.5 in MeOH).

**IR** (neat) 3353 (b), 2937 (w), 2846 (w) 2797 (w), 1622 (m), 1516 (s), 1491 (s), 1460 (m), 1379 (m), 1270 (m), 1249 (m), 1181 (m), 1076 (w), 1051 (m), 943 (w), 836 (m), 742 (m), 707 (m) $cm^{-1}$.

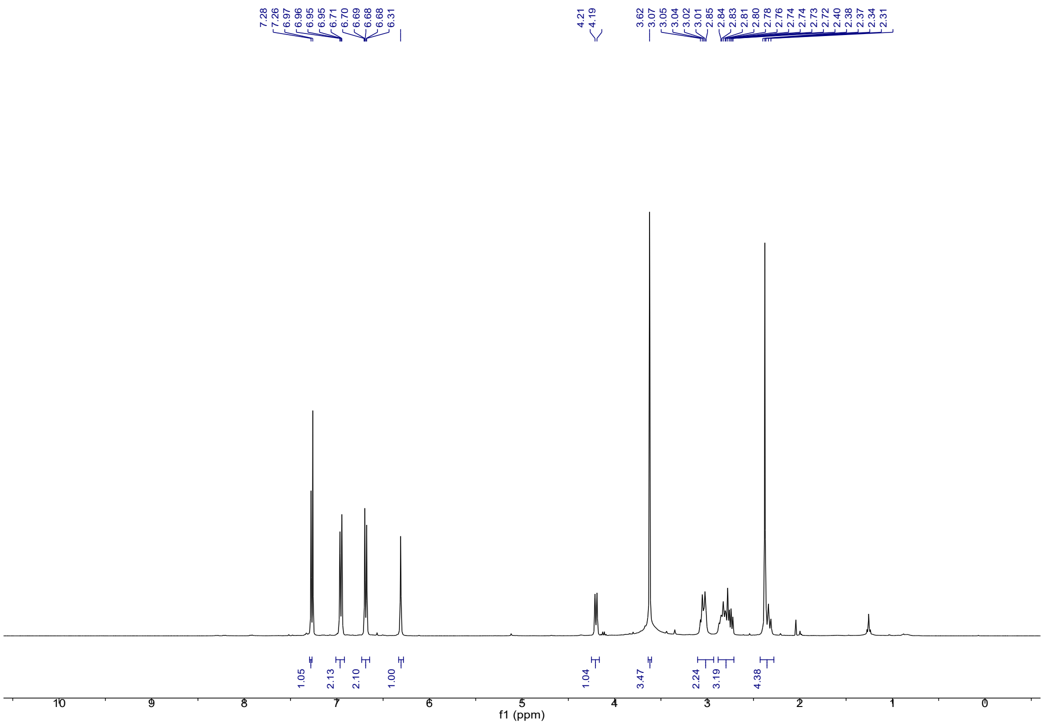
7.28
7.26
6.97
6.96
6.95
6.95
6.71
6.70
6.69
6.68
6.68
6.31
4.21
4.19
3.62
3.07
3.05
3.04
3.02
3.01
2.85
2.84
2.83
2.81
2.80
2.78
2.76
2.74
2.74
2.73
2.72
2.40
2.38
2.37
2.34
2.31
1.05
2.13
2.10
1.00
1.04
3.47
2.24
3.19
4.38
f1 (ppm)

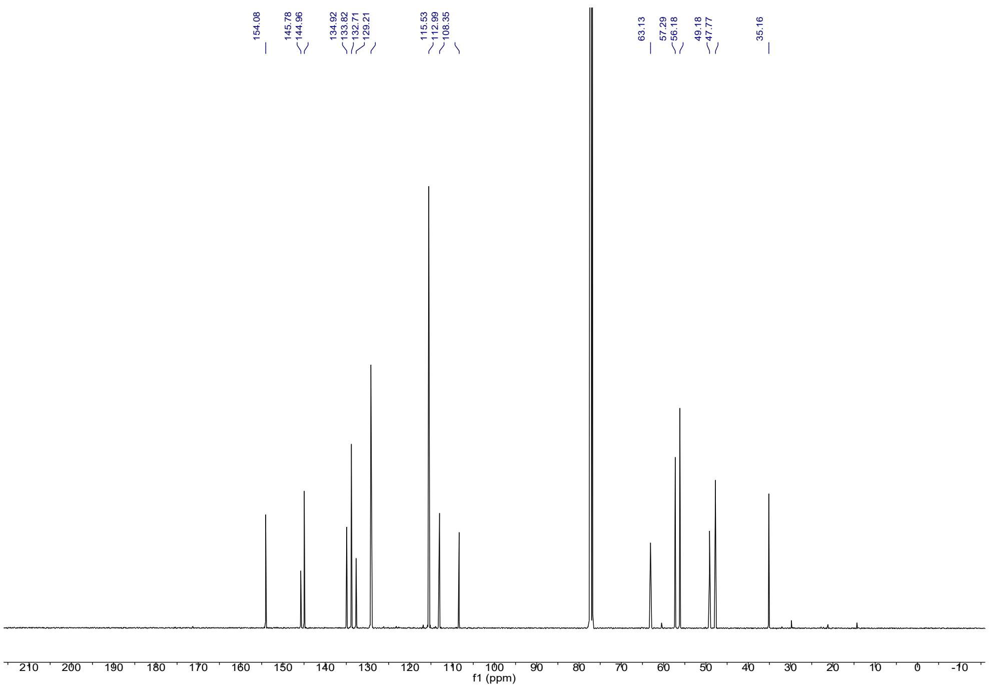
154.08
145.78
144.96
134.92
133.82
132.71
129.21
115.53
112.99
108.35
63.13
57.29
56.18
49.18
47.77
35.16
f1 (ppm)

##### 2.2.7 Azobenzene 10

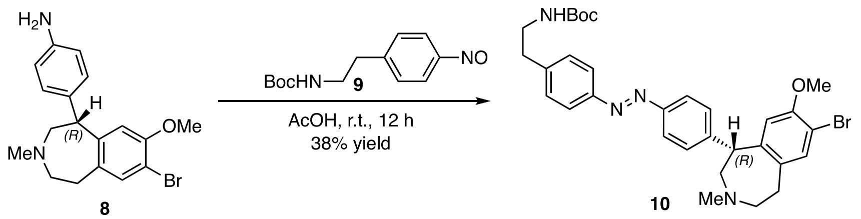


In a 150 mL round bottom flask, *tert*-butyl (4-aminophenethyl)carbamate (0.181 g, 0.767 mmol, 1.50 eq) was dissolved in DCM (13 mL). Oxone (1.57 g, 2.55 mmol, 5.00 eq) and water (13 mL) were added, and the biphasic mixture was vigorously stirred for 3h. The green organic layer was separated off, and the aqueous phase was extracted with DCM (20 mL x 2). The combined organic layers were washed with bicarbonate (sat., aq., 10 mL) and dried over $Na_2SO_4$. The organic phase was filtered and concentrated under reduced pressure to an around 3 mL, green solution containing the intermediate nitrosobenzene (**9**). In a separate 50 mL round bottom flask, **8** (185 mg, 0.511 mmol, 1.00 eq) in AcOH (7.7 mL) and the prepared nitrosobenzene **9** was added as a solution in DCM. The DCM was evaporated at 500 mbar on the rotary evaporator and the reaction mixture stirred for 12h. The reaction mixture was concentrated on the rotary evaporator and the crude oil subjected to column chromatography ($SiO_2$, 24g, 0-10% MeOH in DCM) to yield the product **10** as yellow oil in 38% yield (115 mg, 0.195 mmol).

$\mathbf{R}_f$ = 0.16 (1:1 EA:hexanes; Vis, yellow spot).

**LCMS** (5-100% MeCN in $H_2O$ with 0.1% formic acid over 5 min) $t_R$ = 3.863 min, 360 nm detection.

**LRMS** (ESI): calc. for $C_{31}H_{38}BrN_4O_3^+[M+H]^+$: 593.2122; found 593.2.

**HRMS** (APCI): calc. for $C_{31}H_{38}BrN_4O_3^+[M+H]^+$: 595.2101; found 595.2100.

**$^1$H NMR** (400 MHz, $CDCl_3$) δ 7.9 – 7.8 (m, 4H), 7.4 – 7.3 (m, 5H), 6.3 (s, 1H), 4.4 (d, *J* = 8.1 Hz, 1H), 3.6 (s, 3H), 3.4 – 3.3 (m, 2H), 3.2 – 2.9 (m, 3H), 2.9 – 2.7 (m, 4H), 2.4 – 2.3 (m, 4H), 1.4 (s, 9H).

**$^{13}$C NMR** (101 MHz, $CDCl_3$) δ 155.9, 154.1, 151.3, 151.3, 145.5, 144.2, 142.5, 134.7, 134.0, 129.5, 129.0, 123.1, 123.0, 112.8, 108.8, 79.2, 62.1, 56.9, 56.0, 49.4, 47.4, 41.6, 36.1, 34.6, 28.4.

**IR** (neat) 3331 (b), 2938 (m), 2798 (m), 1598 (m), 1515 (s), 1494 (m), 1461 (w), 1384 (w), 1343 (s), 1251 (m), 1077 (w), 1032 (m), 921 (w), 855 (m), 843 (w), 815 (w), 714 (m) $cm^{-1}$.

**$^{1}$H NMR**

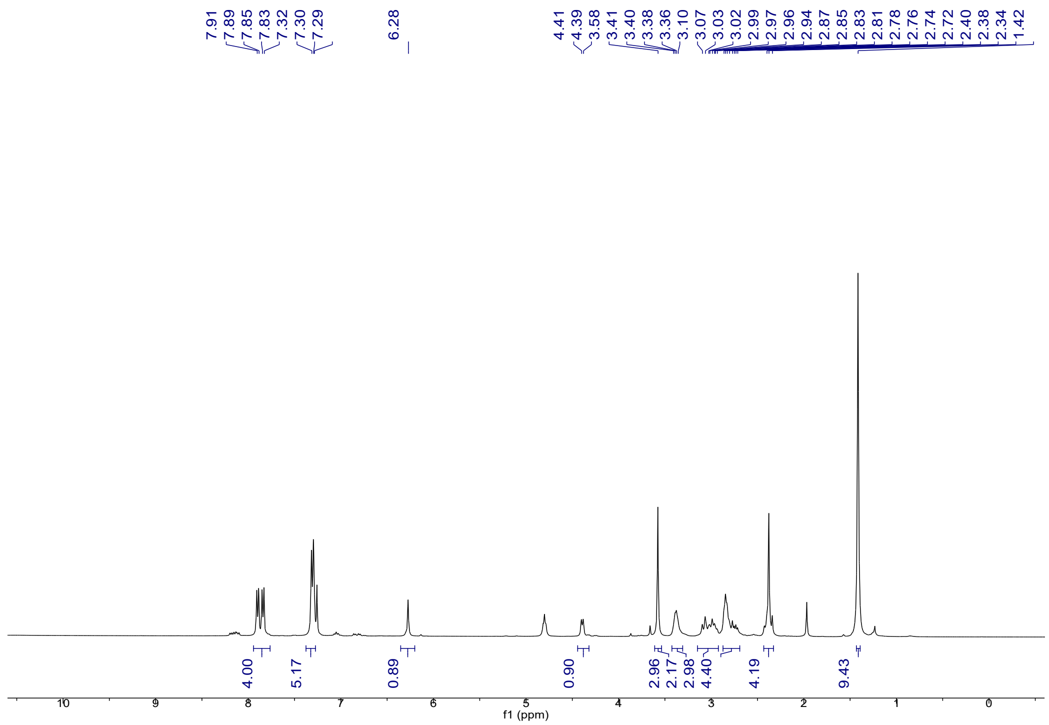


**$^{13}$C NMR**

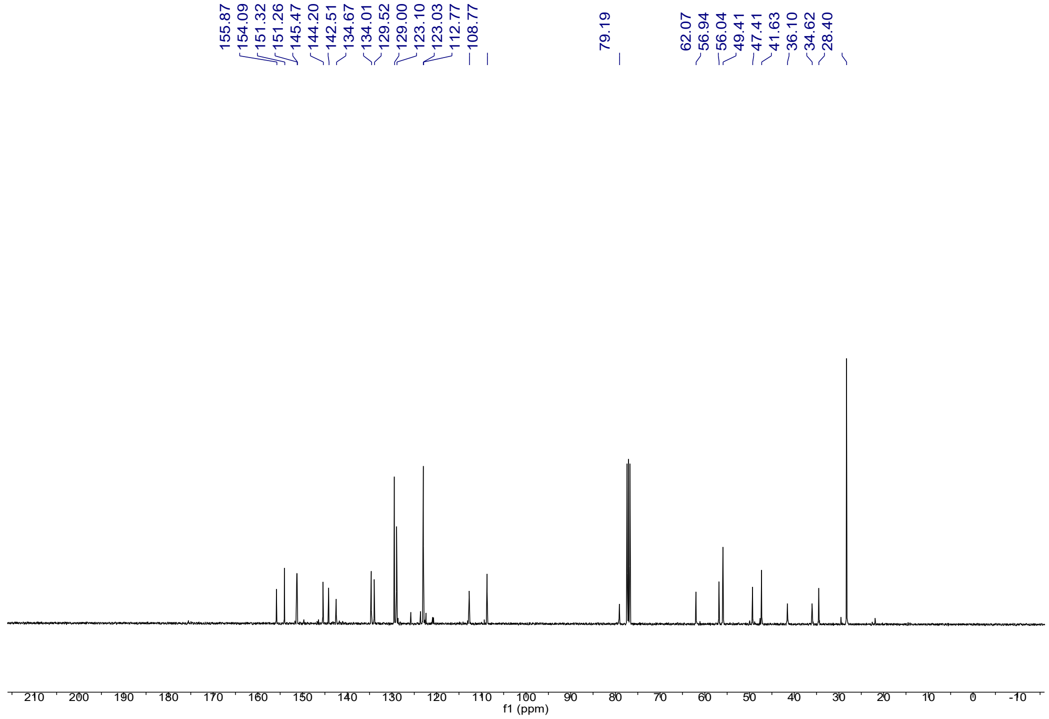

##### 2.2.8 Phenol-Benzazepine Azo 11

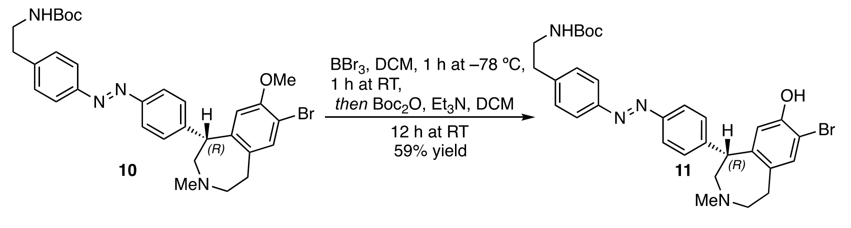

In a flame-dried 100 mL round bottom flask under magnetic stirring, **10** (90.8 mg, 0.153 mmol, 1.00 eq) was dissolved in DCM (22 mL). The solution was cooled to –78 °C in a dry ice/acetone bath. In a separate flame-dried 10 mL round bottom flask, $BBr_3$ (72 µL, 0.765 mmol, 5.00 eq) was dissolved in hexane (3.3 mL). The $BBr_3$-solution was added dropwise to the reaction flask at –78 °C and stirred for 1h. The reaction was then allowed to warm to ambient temperature and continued to stir for 1h. After this time, the reaction was cooled back to –78 °C and MeOH (75 mL) was slowly added to quench remaining $BBr_3$. The mixture was concentrated under reduced pressure, and the crude material co-evaporated with MeOH (2 x 10 mL). To re-protect any deprotected amine, the crude oil was dissolved in DCM (1.5 mL) and $Et_3N$ (213 µL, 1.53 mmol, 10.0 eq) and $Boc_2O$ (40.1 mg, 0.184 mmol, 1.20 eq) were added to the solution. This brown suspension was stirred at ambient temperature for 12h. The reaction mixture was treated with $NaHCO_3$ (aq., 5 mL) and extracted with DCM (3 x 15 mL). The combined organic phases were dried with $Na_2SO_4$, filtered, and concentrated under reduced pressure to yield a crude oil. The crude product was purified via flash column chromatography ($SiO_2$, 12g, 0-10% MeOH in DCM) and the free phenol product **11** was obtained as an orange oil in 59% yield (52.0 mg, 0.090 mmol).

$\mathbf{R}_f$ = 0.34 (5% MeOH and 15% DCM in EtOAc; UV; yellow spot).

**LCMS** (5-100% MeCN in $H_2O$ with 0.1% formic acid over 5 min) $t_R$ = 3.718 min, 254 nm detection.

**LRMS** (ESI): calc. for $C_{30}H_{36}BrN_4O_3^+$ $[M+H]^+$: 579.2; found 579.2.

**HRMS** (APCI): calc. for $C_{30}H_{36}BrN_4O_3^+$ $[M+H]^+$: 581.1945; found 581.1949.

**IR** (neat) 3405 (b), 2916 (w), 1761 (w), 1701 (m), 1600 (m), 1498 (m), 1434 (m), 1405 (m), 1365 (m), 1341 (w), 1309 (w), 1246 (m), 1162 (m), 1014 (S), 950 (m), 853 (w) $cm^{-1}$.

**$^1$H NMR** (400 MHz, $CDCl_3$) δ 7.9 – 7.8 (m, 4H), 7.3 (d, *J* = 8.2 Hz, 2H), 7.2 (s, 1H), 7.2 (d, *J* = 8.2 Hz, 2H), 6.2 (s, 1H), 4.2 (d, *J* = 8.8 Hz, 1H), 3.5 – 3.4 (m, 2H), 3.1 – 3.0 (m, 2H), 3.0 – 2.8 (m, 4H), 2.7 – 2.7 (m, 1H), 2.4 – 2.2 (m, 4H), 1.4 (s, 9H).

**$^{13}$C NMR** (101 MHz, $CDCl_3$) δ 156.0, 151.5, 151.5, 151.4, 145.7, 145.1, 142.6, 134.2, 133.0, 129.7, 129.6, 129.3, 129.2, 129.2, 123.2, 123.2, 116.5, 107.5, 79.5, 62.6, 57.2, 48.9, 47.3, 41.7, 36.2, 34.8, 28.5.

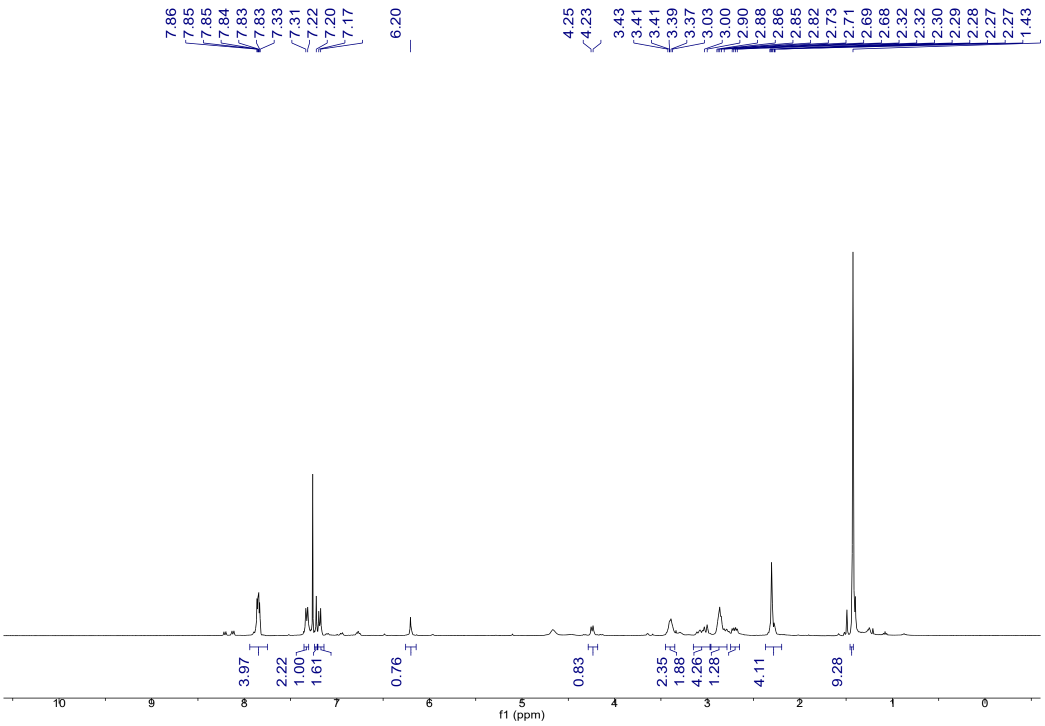
7.86 7.85 7.85 7.84 7.83 7.83 7.33 7.31 7.22 7.20 7.17 6.20
4.25 4.23 3.43 3.41 3.39 3.37 3.03 3.00 2.90 2.88 2.86 2.85 2.82 2.73 2.71 2.69 2.68 2.32 2.32 2.30 2.29 2.28 2.27 2.27 1.43
3.97 2.22 1.00 1.61 0.76 0.83 2.35 1.88 4.26 1.28 4.11 9.28
f1 (ppm)

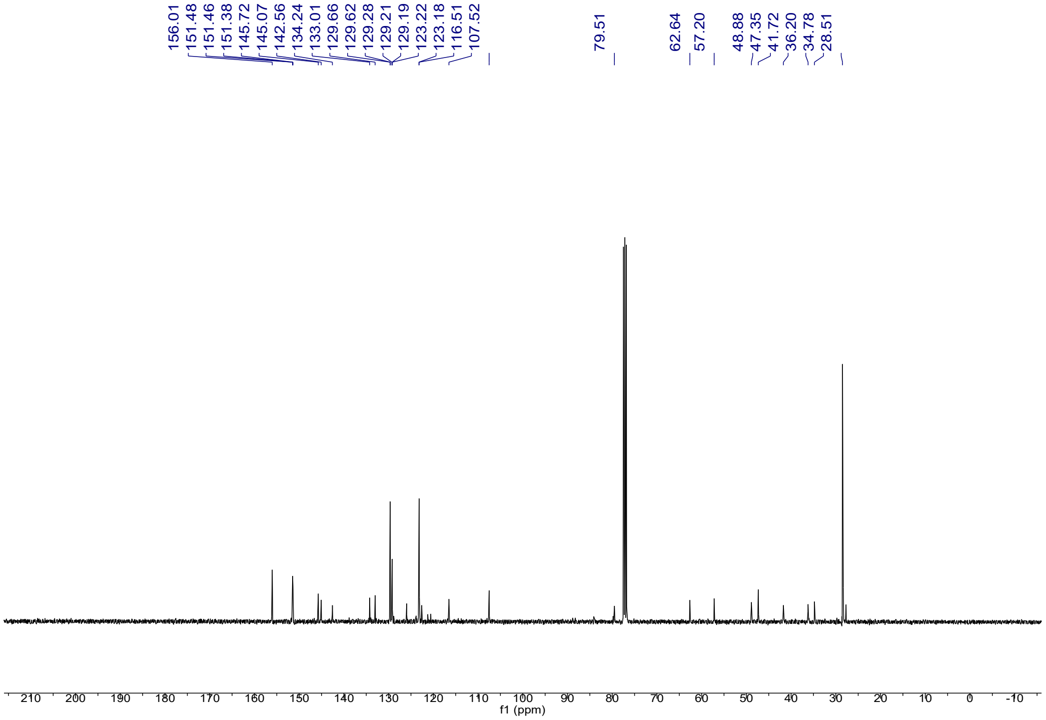
156.01 151.48 151.46 151.38 145.72 145.07 142.56 134.24 133.01 129.66 129.62 129.28 129.21 129.19 123.22 123.18 116.51 107.52
79.51
62.64 57.20
48.88 47.35 41.72 36.20 34.78 28.51
f1 (ppm)

##### 2.2.9 Azo-Peg[12] 12

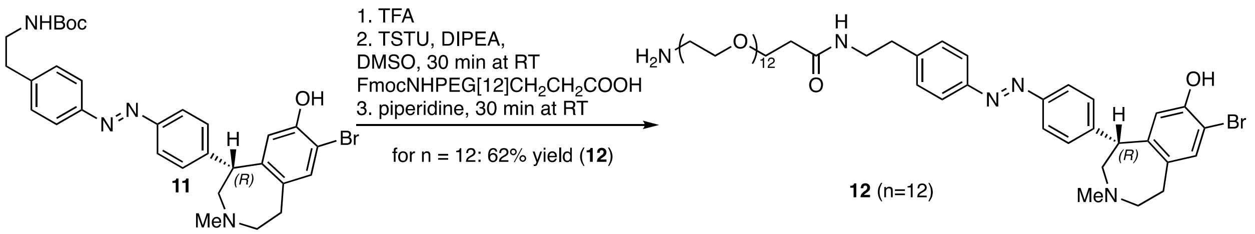


In a 1.5 mL vial under ambient conditions, FmocNHPEG[12]$CH_2CH_2COOH$ (32.1 mg, 0.038 mmol, 2.7 eq) was dissolved in DMSO (109 uL). TSTU (11.5 mg, 0.038 mmol, 2.7 eq) and DIPEA (20 uL, 0.115 mmol, 8.1 eq) were added. The reaction was stirred for 30 min until full conversion was determined by LCMS analysis. In the meanwhile, in a 1.5 mL vial, azobenzene **11** (8.2 mg, 0.014 mmol, 1.0 eq) was Boc-deprotected by treatment with TFA (100 uL). The TFA was removed after 30 seconds by evaporation under a nitrogen stream. The crude azobenzene amine was dissolved in DMSO (80 uL) and added to the activated acid with an additional amount of DIPEA (5 uL, 0.028 mmol, 2.0 eq). The reaction was stirred for 1h, until full conversion was determined by LCMS. Piperidine (140 uL, 1.415 mmol, 100 eq) was added to the reaction mixture and stirring was continued until full conversion was determined by LCMS. The crude reaction mixture was cooled in an ice-water bath and treated with HOAc (97 uL, 1.698 mmol, 120 eq). The mixture was purified by RP-HPLC (15-35% MeCN in $H_2O$ with 0.1% formic acid over 5 min, semi-preparative column, 9.0 mL/min flowrate, detection at 360 nm, $t_R$ = 2.791 min) to yield thePEG-amine product **12** after lyophilization as yellow oil in 62% yield (9.4 mg, 0.009 mmol).

**LCMS** (5-100% MeCN in $H_2O$ with 0.1% formic acid over 5 min) $t_R$ = 2.791 min, 360 nm detection.

**LRMS** (ESI): calc. for $C_{52}H_{82}BrN_5O_{14}^{2+}$ $[M+2H]^{2+}$: 539.75; found 539.8.

**HRMS** (APCI): calc. for $C_{52}H_{81}BrN_5O_{14}^{+}$ $[M+H]^{+}$: 1080.4937; found 1080.4944.

**$^1$H NMR** (400 MHz, MeOD) δ 8.0 (d, $J$ = 8.2 Hz, 2H), 7.9 (d, $J$ = 8.2 Hz, 2H), 7.5 – 7.4 (m, 4H), 7.3 (s, 1H), 6.3 (s, 1H), 4.5 (d, $J$ = 8.6 Hz, 1H), 3.8 – 3.8 (m, 3H), 3.7 – 3.6 (m, 43H), 3.5 (t, $J$ = 7.1 Hz, 3H), 3.3 (s, 1H), 3.2 – 3.1 (m, 5H), 3.0 – 2.8 (m, 3H), 2.7 (t, $J$ = 6.4 Hz, 1H), 2.5 (s, 4H), 2.4 (t, $J$ = 6.0 Hz, 2H).

**$^{13}$C NMR** (101 MHz, MeOD) δ 172.6, 152.5, 151.5, 151.3, 145.4, 143.9, 143.2, 133.4, 132.8, 129.5, 129.1, 129.0, 122.9, 122.6, 116.1, 106.7, 70.1, 70.1, 70.1, 70.0, 70.0, 70.0, 70.0, 69.9, 69.9, 69.9, 69.8, 69.8, 69.8, 69.8, 69.7, 69.7, 69.7, 69.6, 69.5, 69.3, 69.3, 67.1, 66.9, 66.6, 66.5, 61.9, 56.9, 42.6, 40.2, 39.3, 39.3, 34.9, 33.2, 33.0.

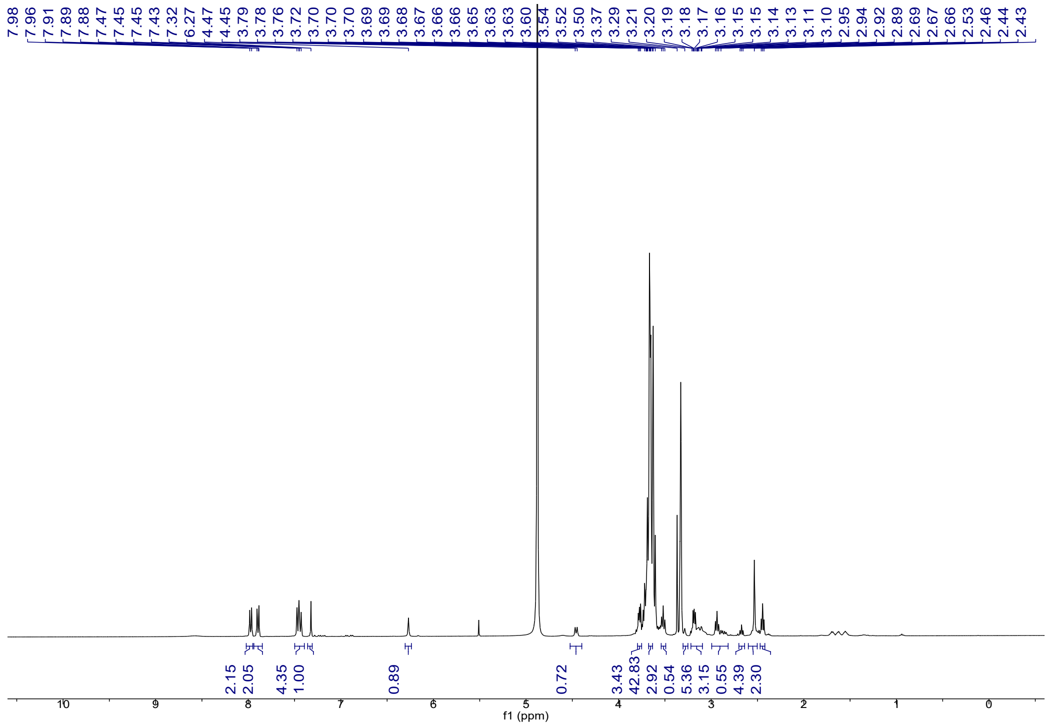


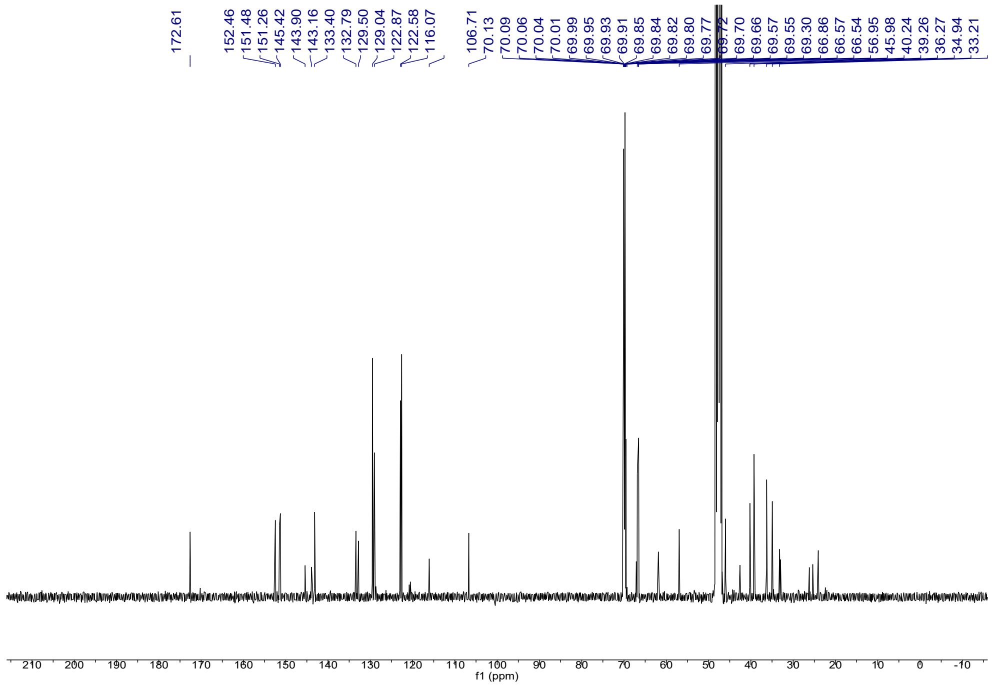

##### 2.2.10 P-D1$_{block12}$

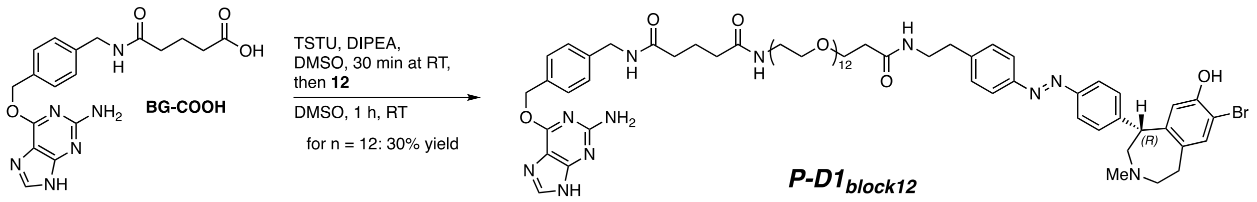

In a 1.5 mL vial, BG-COOH (2.7 mg, 0.007 mmol, 1.5 eq) was dissolved in DMSO (36 uL). TSTU (2.1 mg, 0.007 mmol, 1.5 eq) and DIPEA (3 uL, 0.019 mmol, 4.0 eq) were added and the reaction mixture was stirred for 30 min, until full conversion was determined by LCMS analysis. **12** (5.0 mg, 0.005 mmol, 1.0 eq) was added to the reaction mixture as a solution in DMSO (26 uL) and more DIPEA (3 uL, 0.019 mmol, 4.0 eq) was added. After the reaction was judged complete by LCMS analysis, the mixture was purified by RP-HPLC (10-45% MeCN in $H_2O$ with 0.1% formic acid over 9 min, semi-preparative column, 9.0 mL/min flow rate, detection at 360 nm, $t_R$ = 6.397 min) to yield the product after evaporation of the solvent at 40 °C in vacuo as a yellow oil in 30% yield (2.0 mg, 0.001 mmol). **P-D1$_{block12}$** was synthesized using starting materials of >99% *ee*. Furthermore, a **P-D1$_{block12,S}$** version was synthesized using *S*-configured starting materials of 56% *ee*.

**LCMS** (5-100% MeCN in $H_2O$ with 0.1% formic acid over 5 min) $t_R$ = 2.826 min, 360 nm detection.

**LRMS** (ESI): calc. for $C_{70}H_{100}BrN_{11}O_{17}^{2+}$ $[M+2H]^{2+}$: 722.8236; found 722.8.

**HRMS** (ESI): calc. for $C_{70}H_{100}BrN_{11}O_{17}^{2+}$ $[M+2H]^{2+}$: 722.8236; found 722.8223.

**$^1$H NMR** (400 MHz, MeOD) δ 8.0 (d, *J* = 8.3 Hz, 2H), 7.9 – 7.8 (m, 3H), 7.5 (d, *J* = 7.9 Hz, 2H), 7.4 (t, *J* = 8.6 Hz, 4H), 7.4 (s, 1H), 7.3 (d, *J* = 7.8 Hz, 2H), 6.3 (s, 1H), 5.5 (s, 2H), 4.5 (d, *J* = 8.8 Hz, 1H), 4.4 (s, 2H), 3.7 (t, *J* = 6.0 Hz, 2H), 3.6 – 3.6 (m, 47H), 3.5 – 3.5 (m, 4H), 3.4 – 3.4 (m, 1H), 3.3 – 3.1 (m, 2H), 2.9 (t, *J* = 7.2 Hz, 3H), 2.7 (s, 4H), 2.4 (t, *J* = 6.0 Hz, 2H), 2.3 (dt, *J* = 14.4, 7.4 Hz, 4H), 1.9 (q, *J* = 7.4 Hz, 2H).

**$^{13}$C NMR** (101 MHz, MeOD) δ 175.4, 175.2, 174.1, 161.7, 154.2, 153.0, 152.6, 145.9, 144.7, 140.1, 137.0, 135.0, 130.9, 130.4, 129.7, 128.8, 124.4, 124.0, 117.5, 108.4, 71.6, 71.5, 71.5, 71.4, 71.3, 71.2, 70.5, 68.6, 68.3, 62.7, 58.1, 47.0, 43.9, 41.6, 40.4, 37.7, 36.3, 36.2, 36.1, 33.8, 23.2.

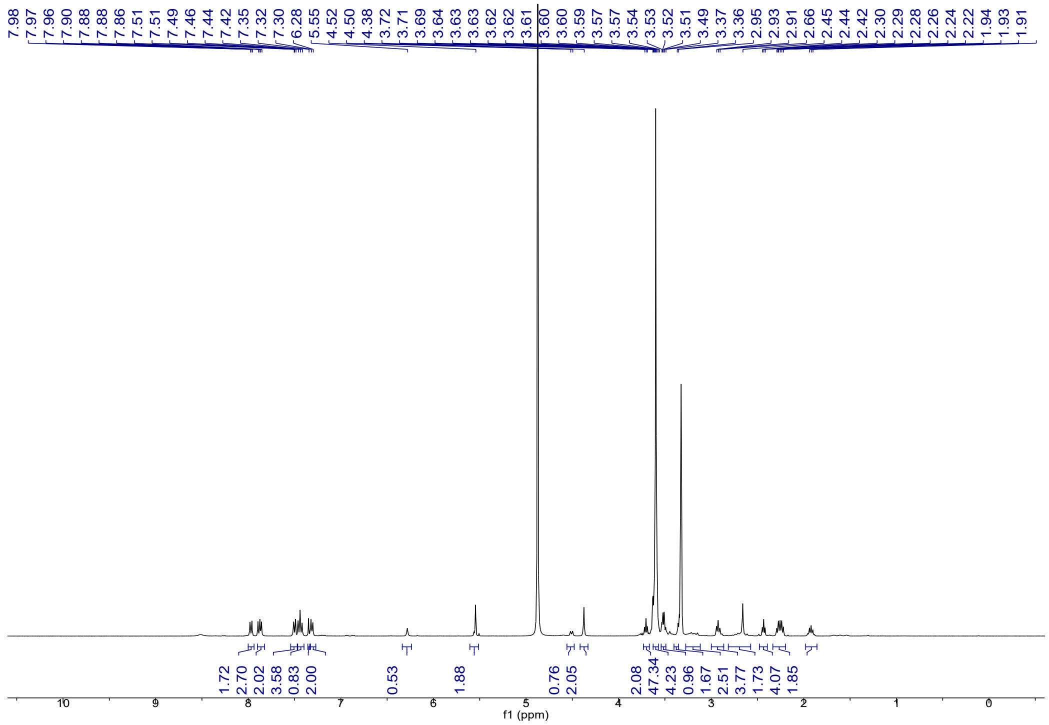

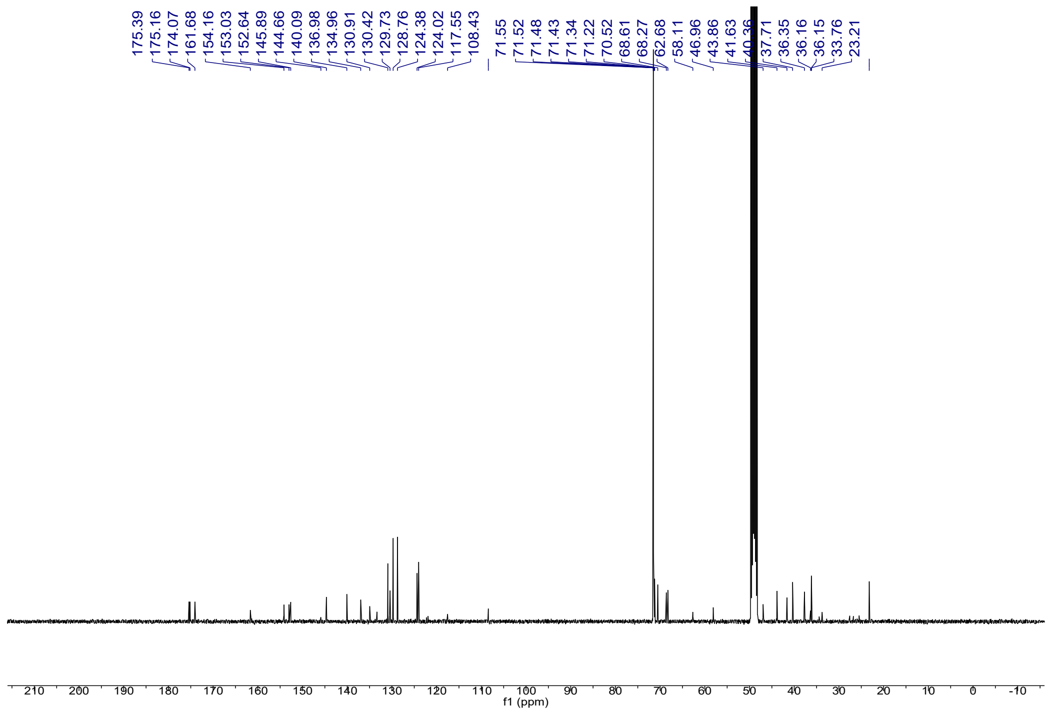

##### 2.2.11 Azo-PEG[24] 13

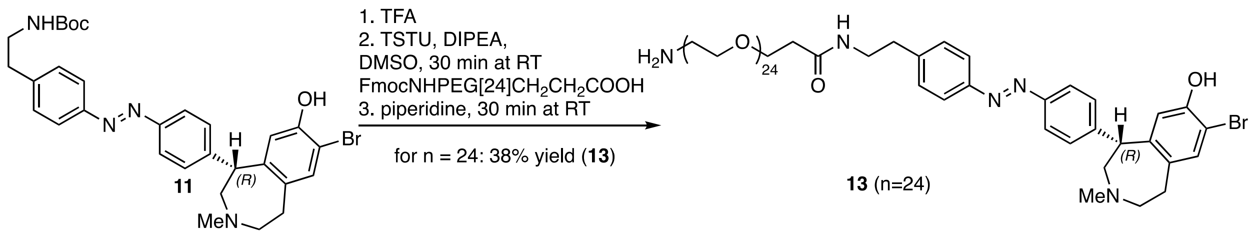

In a 1.5 mL vial under ambient conditions, FmocNHPEG[24]$CH_2CH_2COOH$ (28.3 mg, 0.021 mmol, 1.2 eq) was dissolved in DMSO (133 uL). TSTU (6.2 mg, 0.021 mmol, 1.2 eq) and DIPEA (24 uL, 0.140 mmol, 8.1 eq) were added. The reaction was stirred for 30 min until full conversion was determined by LCMS analysis. In the meanwhile, in a 1.5 mL vial, **11** (10.0 mg, 0.017 mmol, 1.0 eq) was Boc-deprotected by treatment with TFA (100 uL). The TFA was removed after 30 seconds by evaporation under a nitrogen stream. The crude azobenzene-amine was dissolved in DMSO (98 uL) and added to the activated acid with an additional amount of DIPEA (6 uL, 0.035 mmol, 2.0 eq). The reaction was stirred for 1h, until full conversion was determined by LCMS. Piperidine (171 uL, 1.726 mmol, 100 eq) was added to the reaction mixture and stirring was continued until full conversion was determined by LCMS. The crude reaction mixture was cooled in an ice-water bath and treated with HOAc (119 uL, 2.071 mmol, 120 eq). The mixture was purified by RP-HPLC (10-50% MeCN in $H_2O$ with 0.1% formic acid over 6 min, semi-preparative column, 9.0 mL/min flow rate, detection at 360 nm, $t_R$ = 4.300 min) to yield the PEG-amine product **13** after lyophilization as yellow oil in 38% yield (10.6 mg, 0.007 mmol).

**LCMS** (5-100% MeCN in $H_2O$ with 0.1% formic acid over 5 min) $t_R$ = 2.847 min, 360 nm detection.

**LRMS** (ESI): calc. for $C_{76}H_{130}BrN_5O_{26}^{2+}$ $[M+2H]^{2+}$: 803.9; found 804.0.

**HRMS** (ESI): calc. for $C_{76}H_{130}BrN_5O_{26}^{2+}$ $[M+2H]^{2+}$: 803.9088; found 803.9061.

**$^1$H NMR** (400 MHz, MeOD) δ 8.1 – 7.8 (m, 4H), 7.5 (d, *J* = 8.1 Hz, 4H), 7.4 (s, 1H), 6.3 (s, 1H), 4.7 (d, *J* = 9.0 Hz, 1H), 3.8 – 3.8 (m, 2H), 3.7 – 3.6 (m, 96H), 3.6 – 3.5 (m, 3H), 3.4 (s, 1H), 3.2 (t, *J* = 5.1 Hz, 2H), 3.1 – 2.9 (m, 5H), 2.9 (s, 3H), 2.4 (t, *J* = 6.0 Hz, 2H).

**$^{13}$C NMR** (101 MHz, MeOD) δ 174.0, 154.5, 153.2, 152.6, 144.8, 135.1, 132.3, 130.9, 130.5, 124.5, 124.0, 117.6, 108.8, 71.5, 71.5, 71.5, 71.4, 71.3, 71.3, 71.3, 71.3, 71.2, 71.2, 71.1, 71.0, 70.8, 70.5, 68.5, 68.2, 67.9, 61.7, 57.7, 47.2, 46.3, 41.6, 40.7, 40.2, 37.7, 36.3, 32.4.

**IR** (neat) 3462 (b), 2918 (m), 2872 (m), 1596 (m), 1457 (w), 1348 (w), 1249 (w). 1103 (s), 951 (w) cm$^{-1}$.

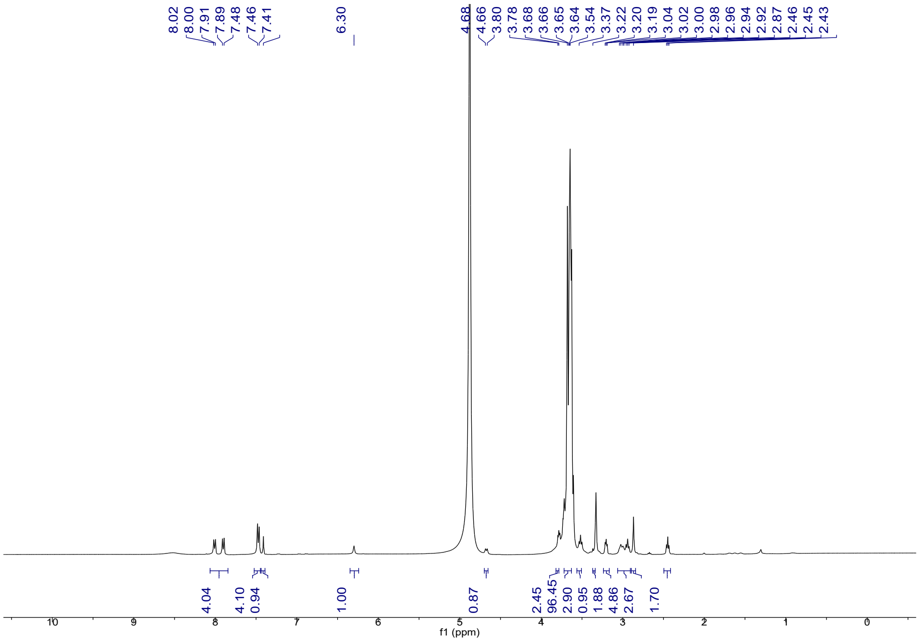


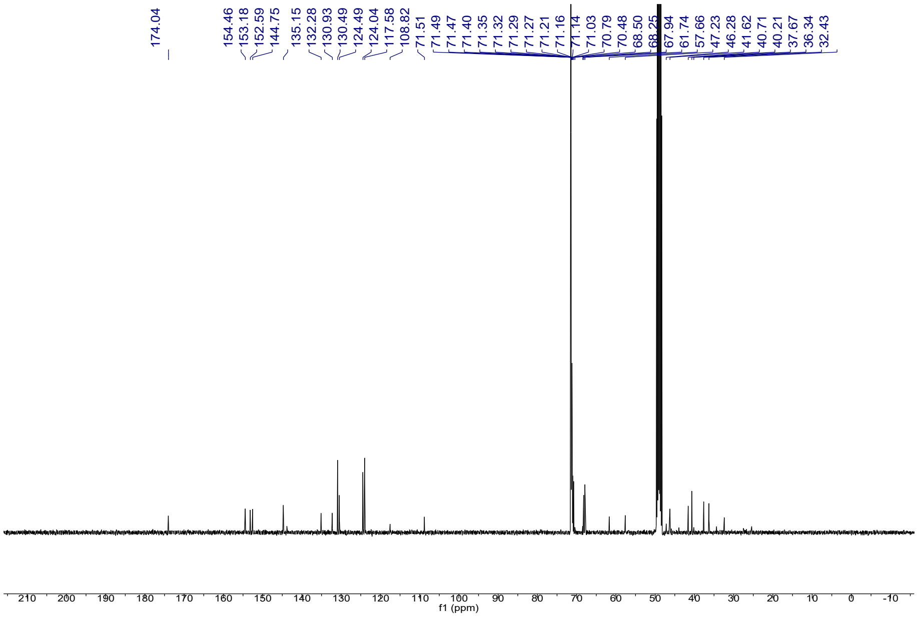

##### 2.2.12 P-D1$_{block24}$

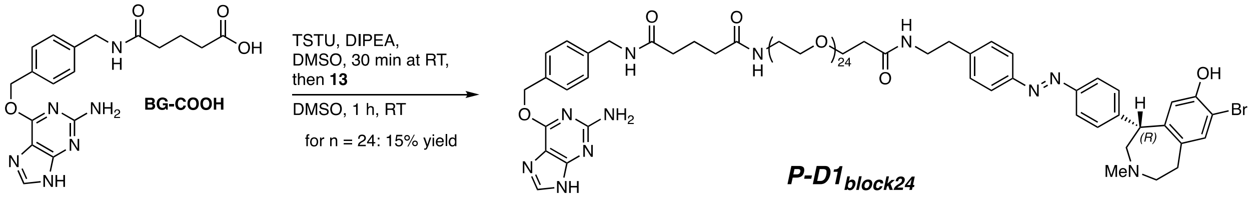


In a 1.5 mL vial, BG-COOH (3.8 mg, 9.9 umol, 1.5 eq) was dissolved in DMSO (36 uL). TSTU (3.0 mg, 9.9 umol, 1.5 eq) and DIPEA (5 uL, 26.4 umol, 4.0 eq) were added and the reaction mixture was stirred for 30 min, until full conversion was determined by LCMS analysis. **13** (10.6 mg, 6.6 umol, 1.0 eq) was added to the reaction mixture as a solution in DMSO (51 uL) and more DIPEA (2 uL, 13.2 umol, 2.0 eq) was added. After the reaction was judged complete by LCMS analysis, the mixture was purified by RP-HPLC (10-40% MeCN in $H_2O$ with 0.1% formic acid over 9 min, semi-preparative column, 9.0 mL/min flow rate, detection at 360 nm, $t_R$ = 8.095 min) to yield the product after evaporation of the solvent at 40 °C in vacuo as a yellow oil in 15% yield (1.9 mg, 0.962 umol). **P-D1$_{block24}$** was synthesized using starting materials of >99% *ee*.

**LCMS** (5-100% MeCN in $H_2O$ with 0.1% formic acid over 5 min) $t_R$ = 3.082 min, 360 nm detection.

**LRMS** (ESI): calc. for $C_{94}H_{146}BrN_{11}Na_2O_{29}^{2+}$ $[M+2Na]^{2+}$: 1009.5; found 1009.4.

**HRMS** (ESI): calc. for $C_{94}H_{148}BrN_{11}O_{29}^{2+}$ $[M+2H]^{2+}$: 986.9808; found 986.9844.

**$^1$H NMR** (400 MHz, MeOD) δ 8.0 – 7.9 (m, 5H), 7.5 – 7.4 (m, 6H), 7.4 – 7.3 (m, 3H), 6.3 (s, 1H), 5.6 (s, 2H), 4.4 (d, *J* = 8.7 Hz, 1H), 4.4 (s, 2H), 3.7 (d, *J* = 6.0 Hz, 2H), 3.6 – 3.6 (m, 92H), 3.5 – 3.5 (m, 4H), 3.5 – 3.4 (m, 2H), 3.3 – 3.2 (m, 1H), 3.2 – 3.0 (m, 3H), 2.9 (t, *J* = 7.1 Hz, 2H), 2.9 – 2.8 (m, 1H), 2.5 – 2.4 (m, 6H), 2.3 (dt, *J* = 13.5, 7.5 Hz, 4H), 2.0 – 1.9 (m, 2H).

**$^{13}$C NMR** (101 MHz, MeOD) δ 175.4, 175.2, 174.1, 161.6, 153.8, 152.8, 152.6, 146.9, 145.4, 144.6, 140.1, 137.0, 134.8, 134.3, 130.9, 130.5, 129.7, 128.8, 124.3, 124.0, 117.5, 108.0, 71.5, 71.5, 71.4, 71.3, 71.2, 70.5, 68.6, 68.3, 63.4, 58.4, 49.5, 47.5, 43.9, 41.6, 40.3, 37.7, 36.3, 36.2, 36.2, 34.8, 23.2.

**IR** (neat) 2871 (m), 1738 (w), 1625 (m), 1581 (m), 1455 (w), 1352 (m), 1281 (w), 1104 (s), 944 (m) cm$^{-1}$.

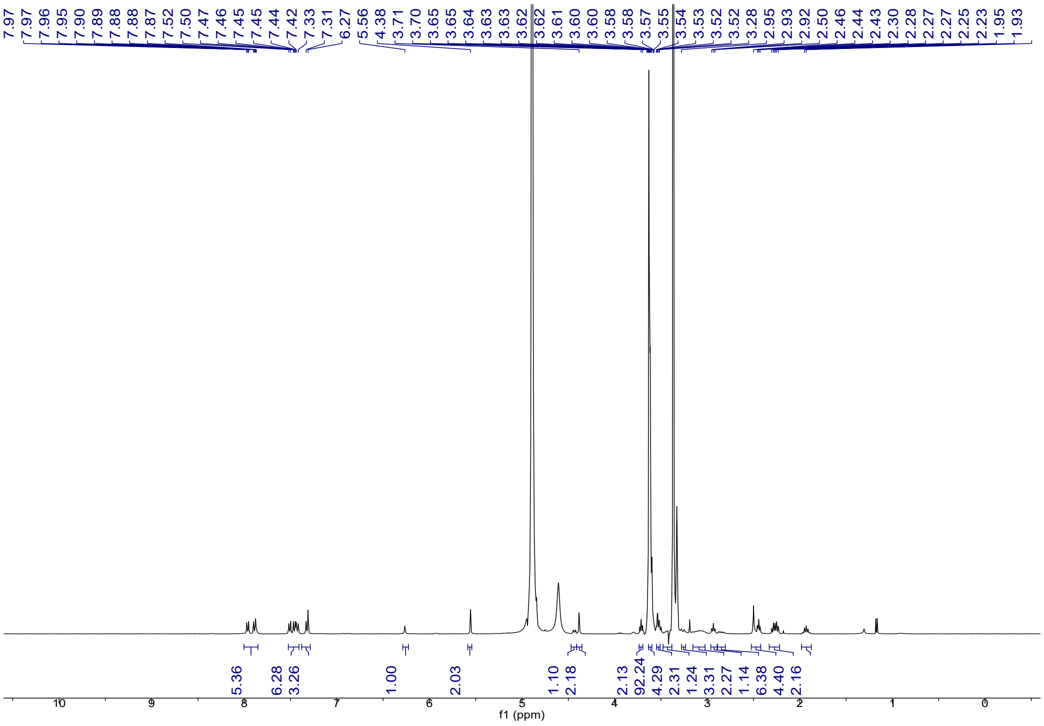
7.97 7.97 7.96 7.95 7.90 7.89 7.88 7.87 7.52 7.50 7.47 7.46 7.45 7.45 7.44 7.42 7.33 7.31 6.27 5.56 4.38 3.71 3.70 3.65 3.64 3.63 3.63 3.62 3.62 3.61 3.60 3.60 3.58 3.58 3.57 3.55 3.54 3.53 3.52 3.52 3.28 2.95 2.93 2.92 2.50 2.46 2.44 2.43 2.30 2.28 2.27 2.27 2.25 2.23 1.95 1.93
5.36 6.28 3.26 1.00 2.03 1.10 2.18 2.13 92.24 4.29 2.31 1.24 3.31 2.27 1.14 6.38 4.40 2.16
f1 (ppm)

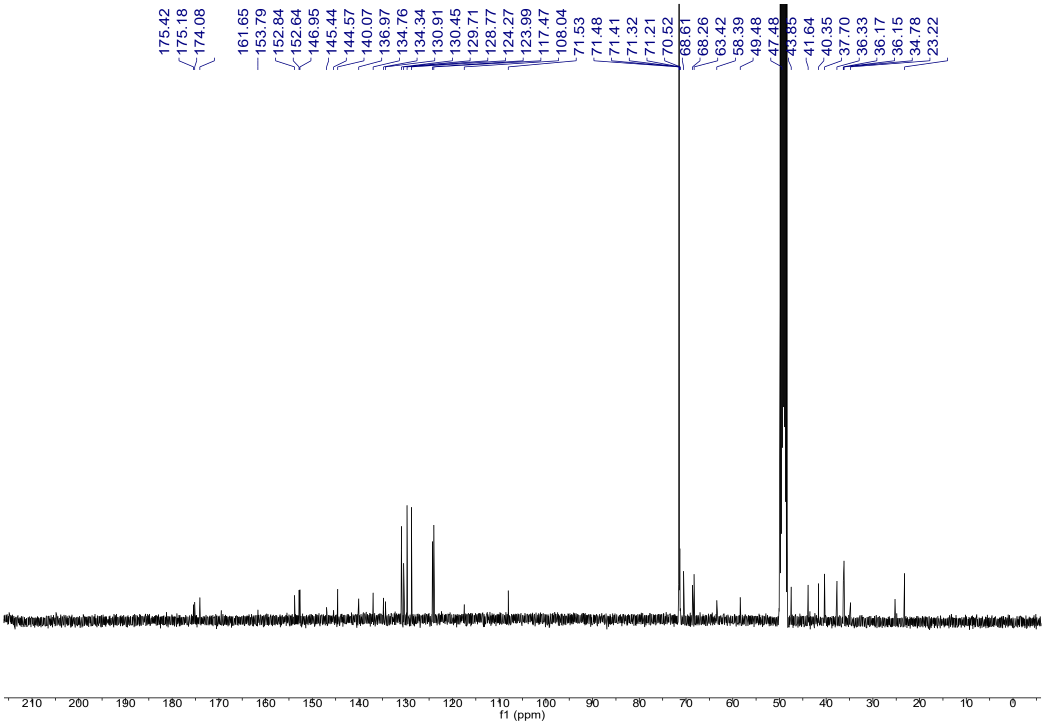
175.42 175.18 174.08 161.65 153.79 152.84 152.64 146.95 145.44 144.57 140.07 136.97 134.76 134.34 130.91 130.45 129.71 128.77 124.27 123.99 117.47 108.04 71.53 71.48 71.41 71.32 71.21 70.52 68.61 68.26 63.42 58.39 49.48 47.48 43.85 41.64 40.35 37.70 36.33 36.17 36.15 34.78 23.22
f1 (ppm)
